# Single-cell multiplex chromatin and RNA interactions in aging human brain

**DOI:** 10.1101/2023.06.28.546457

**Authors:** Xingzhao Wen, Zhifei Luo, Wenxin Zhao, Riccardo Calandrelli, Tri C. Nguyen, Xueyi Wan, John Lalith Charles Richard, Sheng Zhong

## Abstract

**SUMMARY PARAGRAPH:** The dynamically organized chromatin complexes often involve multiplex chromatin interactions and sometimes chromatin-associated RNA (caRNA) ^1–3^. Chromatin complex compositions change during cellular differentiation and aging, and are expected to be highly heterogeneous among terminally differentiated single cells ^4–7^. Here we introduce the <u>Mu</u>lti-Nucleic Acid Interaction Mapping in <u>Si</u>ngle <u>C</u>ell (MUSIC) technique for concurrent profiling of multiplex chromatin interactions, gene expression, and RNA-chromatin associations within individual nuclei. Applied to 14 human frontal cortex samples from elderly donors, MUSIC delineates diverse cortical cell types and states. We observed the nuclei exhibiting fewer short-range chromatin interactions are correlated with an “older” transcriptomic signature and with Alzheimer’s pathology. Furthermore, the cell type exhibiting chromatin contacts between cis expression quantitative trait loci (cis eQTLs) and a promoter tends to be the cell type where these cis eQTLs specifically affect their target gene’s expression. Additionally, the female cortical cells exhibit highly heterogeneous interactions between the XIST non-coding RNA and Chromosome X, along with diverse spatial organizations of the X chromosomes. MUSIC presents a potent tool for exploring chromatin architecture and transcription at cellular resolution in complex tissues.

---

The three-dimensional (3D) folding of the genome is known to exhibit dynamic changes during cellular differentiation processes and demonstrates heterogeneity among terminally differentiated single cells ^4–7^. While its regulatory role in the expression of specific genes has been well-established ^8–10^, the extent to which the 3D genome structure impacts the expression of most genes remains a topic of debate ^11^. Given the pronounced heterogeneity observed in chromatin structure and gene expression across individual cells ^12–14^, a comprehensive understanding of the relationship between 3D genome structure and gene expression at the single-cell resolution is necessary. Therefore, developing single-cell multimodal technologies capable of simultaneously profiling chromatin conformation and gene expression is instrumental for elucidating these intricate relationships.

Single-cell multi-omic technologies enabled joint analysis of chromatin conformation and gene expression ^15–19^ (Supplementary Table 1, Supplemental Note 1). Despite these technical advancements, the simultaneous profiling of multiplex chromatin interactions (co-complexed DNA sequences), gene expression, and RNA-chromatin associations from a single cell remain challenging. To fill this gap, we developed the <u>Mu</u>lti-Nucleic Acid Interaction Mapping in <u>Si</u>ngle <u>C</u>ell (MUSIC) technique, which enables the simultaneous profiling of gene expression and co-complexed DNA sequences with or without co-complexed RNA at the single-cell level.

The architecture of chromatin can encompass both pairwise and multiplex chromatin interactions, highlighting the intricate nature of chromatin complexes ^12,20–22^. ChIA-Drop has facilitated the mapping of multiplex chromatin interactions at single-complex resolution from bulk cells, revealing that multiplex chromatin interactions are prevalent in *Drosophila* ^20^. The MUSIC technique expands the capability to evaluate the composition of pairwise and multiplex chromatin interactions in individual human cells at single-cell resolution.

In addition to DNA, chromatin complexes can also encompass RNA molecules, introducing another layer of complexity to chromatin architecture ^1,2,23^. Chromatin-associated RNA (caRNA) has been shown to contribute to gene expression regulation ^1,24,25^. For instance, the accumulation of the XIST long noncoding RNA (lncRNA) on the X chromosome (XIST-chrX association) is crucial for the silencing of one of the two X chromosomes in female cells, a process known as X-chromosome inactivation (XCI). Various human tissues exhibit both shared and tissue-specific incomplete XCI genes, which are expressed from the silenced X chromosome ^26^. Genes with incomplete XCI can display higher expression levels in females, potentially contributing to sex differences in disease susceptibility ^27^. Recent advancements have enabled genome-wide mapping of RNA-chromatin associations in bulk cells ^1,28–32^. With the application of MUSIC, we can now obtain RNA-chromatin association maps at the single-cell level. Utilizing MUSIC, we uncover cellular heterogeneity in <u>X</u>IST-chrX <u>a</u>ssociation levels (XAL) within the female cortex and explore the co-variation between XAL and chromatin interactions among the female cortical cells.

## Design and workflow

The development of the MUSIC technology is guided by three specific design goals, which collectively enable the joint profiling of gene expression, co-complexed DNA sequences, and RNA-chromatin associations from the same nucleus (Figure 1a). The first goal is to construct RNA and fragmented DNA into a single sequencing library and identify which RNA and DNA sequences originated from the same nucleus. This goal is achieved by labeling all the RNA and the fragmented DNA in the same nucleus with a unique Cell Barcode. This Cell Barcode enables the identification and matching of RNA and DNA sequences originating from the same nuclei.

**Figure 1.**
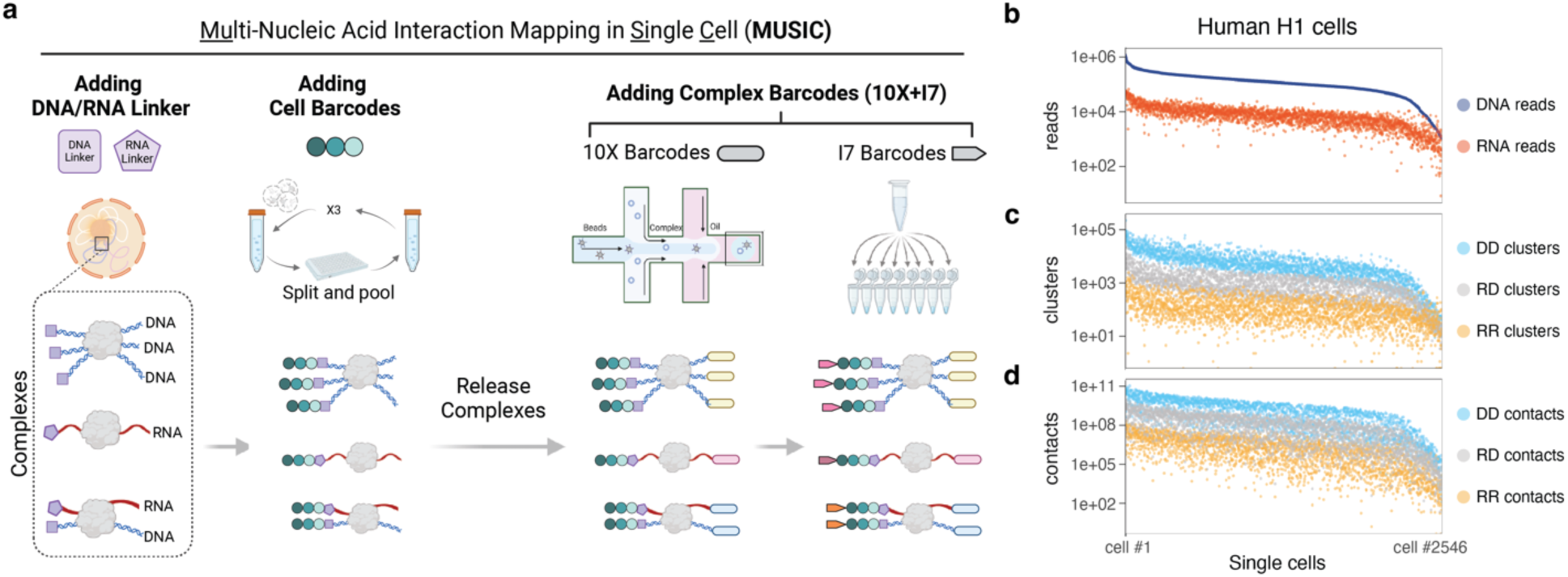
MUSIC workflow and statistics. (a) Schematic view of MUSIC experimental pipeline. (b-d) Summary of MUSIC data in H1 cells. The numbers of uniquely mapped non-duplicate reads (b), clusters (c), and pairwise contacts (d) in every H1 cell (column, n=2,546). DNA-DNA (DD, blue), RNA-DNA (RD, gray), and RNA-RNA (RR, yellow) clusters are separately counted. Multiplex interactions are projected to pairwise interactions and the numbers of pairwise contacts are reported in (d).

The second goal is distinguishing RNA inserts from DNA inserts in the sequencing library. To achieve this, distinct nucleotide sequences are used for the RNA Linker and the DNA Linker, which are ligated to the RNA and DNA molecules, respectively. These linkers are sequenced alongside the RNA and DNA inserts, enabling the differentiation of RNA and DNA molecules within the sequencing data. The third goal is to capture and identify DNA-DNA and RNA-DNA associations, including multi-way contacts. To achieve this, each molecular complex is labeled with a unique Complex Barcode. A molecular complex can encompass various combinations of DNA and RNA, including (1) an isolated RNA molecule, (2) an isolated DNA fragment, (3) multiple DNA fragments, (4) multiple RNA molecules, and (5) at least one RNA molecule and at least one DNA fragment. The Complex Barcodes together with the Cell Barcodes allow for the identification of co-complexed DNA and/or RNA in each cell.

The MUSIC workflow contains two major steps (Supplemental Note 2). The first step ligates *the RNA Linker* to the RNA molecules and *the DNA Linker* to the fragmented DNA and adds the Cell Barcodes (Extended Data Fig.1a-e). The second step adds a Complex Barcode to any RNA or DNA within the same molecular complex. The Complex Barcode consists of a 10X Barcode and a I7 Barcode (Extended Data Fig.1f, g). The final sequencing library is sequenced with a 28bp Read1 sequence, an 8bp index sequence that is the I7 Barcode, and a 150bp Read2 sequence (Extended Data Fig 1h). The 28bp Read1 corresponds to the 10X Barcode, which consists of a 16bp 10X GEM Barcode and a 12bp 10X UMI. Read2 contains the 3rd, 2nd, and 1st Cell Barcodes, the RNA Linker sequence or the DNA-Linker(bt) sequence, and the RNA insert or the DNA insert. It should be noted that each read pair is designed to capture only one insert, either an RNA or a DNA insert, as the Read1 is dedicated to reading the 10X Barcode. This design differs from several ligation-based methods such as Hi-C ^33^ and iMARGI ^28,29^ where each read pair represents two inserts.

## RNA and chromatin interactions in ESC

We applied MUSIC to analyze a mixed population of H1 human and E14 mouse embryonic stem cells (ESC). The resulting mixed-species MUSIC library was sequenced on a NovaSeq platform, generating 3,067,956,666 read pairs. These read pairs resolved 533,233,368 <u>u</u>niquely <u>m</u>apped, <u>n</u>on-<u>d</u>uplicate, and <u>b</u>arcode-<u>c</u>omplete (containing Cell Barcode, 10X Barcode, I7 Barcode, and either DNA Linker or RNA Linker) (UMNDBC) read pairs (Supplementary Table 3). Because as per the experimental design, each UMNDBC read pair contains only one DNA or RNA insert, we will refer to a UMNDBC read pair as a DNA read or an RNA read. This mix-species dataset revealed small species-mixing rates at the cell level and at the chromatin-complex level, supporting the ability of MUSIC to generate data at single-cell and single-complex resolutions (Extended Data Fig. 3a, b, Supplemental Note 3).

We identified 2,546 human H1 cells from this dataset. Each H1 cell contained an average of 144,049 UMNDBC DNA reads, 11,384 UMNDBC RNA reads, corresponding to 7,036 DNA-only clusters (DD clusters), 232 RNA-only clusters (RR clusters), and 1,170 RNA-DNA clusters (RD clusters) (Figure 1b, c), which based on an established procedure ^12^ can be resolved into 2,639,302,084 co-complexed DNA-DNA pairs, 7,089,720 co-complexed RNA-RNA pairs, 250,525,581 co-complexed RNA-DNA pairs (Figure 1d). In total, there were 18,144,410 non-singleton DD clusters, accounting for 55,670,578 DNA reads; 324,121 RR clusters, accounting for 835,184 RNA reads, and 2,401,392 RD clusters, accounting for 13,151,716 RNA reads and 216,515,595 DNA reads (Extended Data Fig. 3c). Among the non-singleton DD clusters, 13,111,228 (72.26%) contained two DNA reads that correspond to pairwise interactions, and 5,033,182 (27.74%) contained three or more DNA reads that respond to multiplex interactions (Extended Data Fig. 3d, Extended Data Fig. 4). In the RD clusters, 1,009,706 (42.05%) contained two reads, i.e., one DNA read and one RNA read, 783,709 (32.64%) contained 3-10 reads, and 607,977 (25.32%) contained more than 10 reads.

We compared MUSIC’s ensemble DNA reads with Micro-C data that were generated from the same human H1 cell line and cultured under the same standard operating protocol as recommended by the 4D Nucleome Consortium. The contact map of MUSIC DD clusters reproduced the structures observed in the Micro-C derived contact map (4DN data portal: 4DNFI2TK7L2F ^34^) (Figure 2a), resulting in a similar distribution of the compartment scores across the genome (Extended Data Fig. 5a). We compared the different-sized MUSIC DD clusters (lower triangles, Figure 2b-d) keeping the same Micro-C dataset as a reference (upper triangles, Figure 2b-d). At the Topologically Associating Domain (TAD) level, MUSIC’s small DD clusters (2-10 DNA reads per cluster) primarily contained contacts within TADs (50 Kb resolution) (Figure 2b). MUSIC’s middle-sized (11-50 reads per cluster) and large clusters (51-100 reads per cluster) recapitulate the TADs and the nested TAD structure (Figures 2c, d). Large clusters revealed more contacts between the nested TADs within a larger TAD (Arrows, Figure 2d).

**Figure 2.**
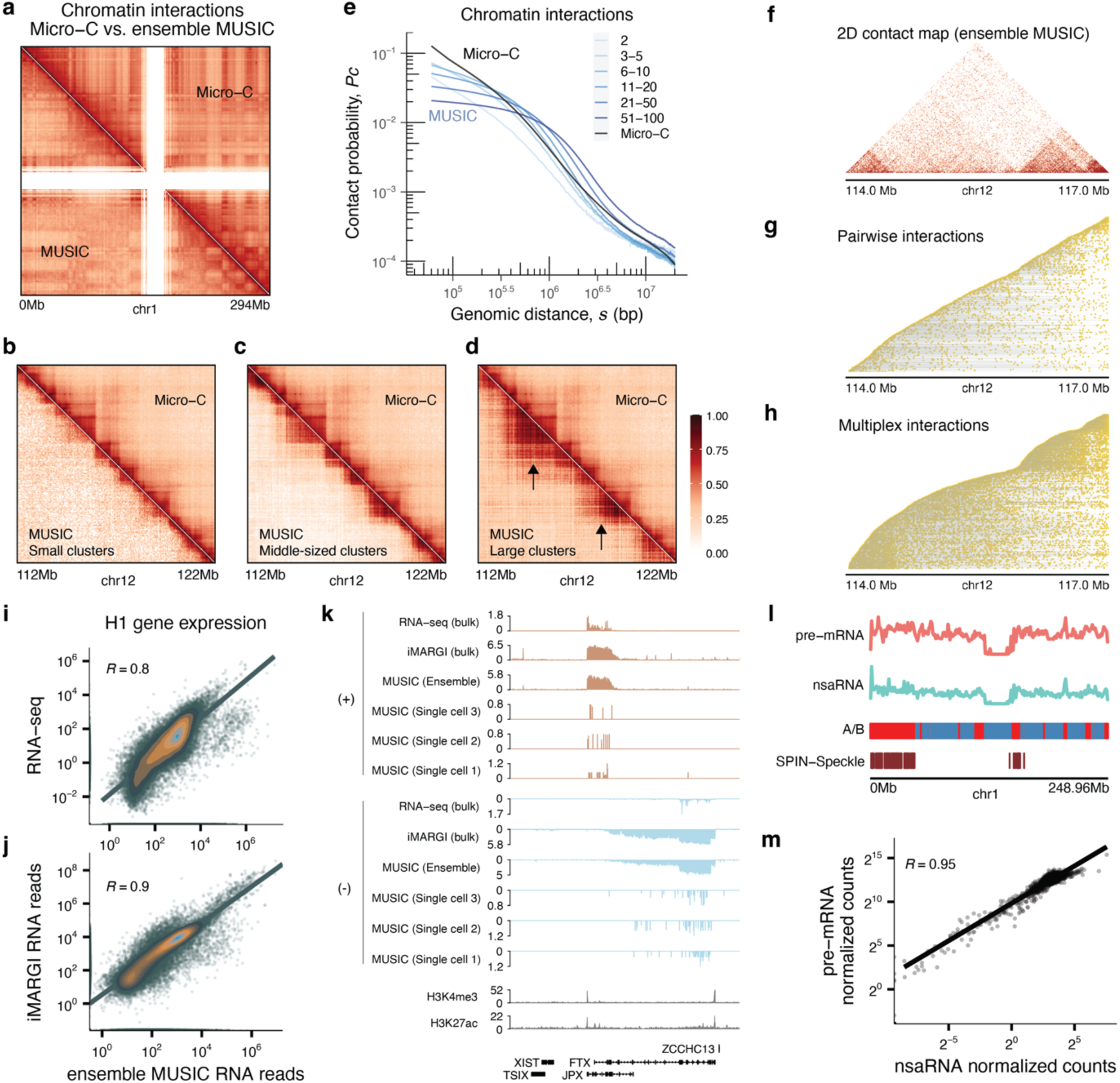
H1 cells MUSIC data. (a) Comparison of Micro-C (upper triangle) and ensemble MUSIC (lower triangle) derived chromatin contact maps on Chromosome 1 at 1Mb resolution. (b-d) Chromatin contact maps based on the ensemble of small (b), middle-sized (c), and large (d) clusters. Resolution: 50Kb. Arrows: Contacts between nested TADs. (e) Pc(s) curves showing the frequency of chromatin contacts (Pc, y axis) vs. genomic distance (s, x axis) for MUSIC DD clusters with different sizes (shades of blue) and Micro-C (black). DNA cluster size: the number of DNA reads in a DD cluster. (f-h) Comparison of 2-D contact map from ensemble MUSIC data (f) with stacked maps of distinct DNA-DNA clusters. Each row represents a cluster, ordered by the smallest genomic coordinate of any DNA read. Yellow dots denote genomic locations of DNA reads within a cluster. Clusters with 2 DNA reads (pairwise interactions) are shown in (g), while clusters with 3 or more DNA reads (multiplex interactions) are shown in (h). (i-j) Scatterplots of RNA levels measured by Reads Per Kilobase (RPK) for every gene (dot) in ensemble MUSIC (x axis) vs. RNA-seq (y axis) (i) and iMARGI (y axis) (j). R: Spearman correlation. (k) RNA reads from RNA-seq, iMARGI, and MUSIC mapped to both strands (+ and -) in human H1 cells. Ensemble MUSIC and individual MUSIC data from three single cells are presented. (l) Distribution of chromatin-associated pre-mRNA and nsaRNA on Chromosome 1 as measured by ensemble MUSIC in H1 cells. Micro-C derived A/B compartments are colored in red/blue. The “Speckle compartmentalization” derived from the SPIN model is denoted in the SPIN-Speckle track. (m) Scatter plot of normalized counts of pre-mRNA reads (y axis) and nsaRNA reads (x axis) in every 1 Mb genomic bin (dot) across the entire genome, based on Ensemble MUSIC data in H1 cells.

To visualize the clusters, we plotted each cluster in a row, with every DNA read of this cluster aligned to their respective genomic coordinates. We ordered the clusters by the genomic coordinates of their leftmost DNA reads. This way, we created a stacked map of the clusters (Figure 2g, h). By comparing the 2-D contact map based on ensemble MUSIC data (Figure 2f) to the stacked maps, we observed that pairwise interactions (clusters with two DNA reads) alone poorly reflected the TAD structure (Figure 2g), while multiplex interactions (clusters with three or more DNA reads) recapitulated the TAD structure (Figure 2h). An analysis involving downsampling the reads suggested that this difference was not due to the different read numbers in pairwise and multiplex interactions (Supplemental Note 4, Extended Data Fig. 5b). This analysis corroborates the difference in the contact maps of different cluster sizes (Figure 2b-d) to suggest that the TAD chromatin structure predominantly consists of multiplex chromatin interactions. Additionally, compared to the pairwise contacts, the multiplex interactions displayed higher contact frequencies at submegabase to several megabase genomic distances, indicating enrichment of long-range chromatin interactions in the multiplex complexes (Figure 2e, Supplemental Note 5).

We compared MUSIC’s ensemble RNA reads (RNA ensemble) from the 2,546 H1 cells to RNA measurements obtained from two bulk assays in H1. Using all 60,719 genes defined in GENCODE v36, we quantified the RNA level of each gene in terms of reads <u>p</u>er <u>k</u>ilobase (RPK). The RPKs of MUSIC’s RNA ensemble correlated with those of bulk RNA-seq (ENCSR000COU ^35^) (Figure 2i, rho = 0.8, p-value = 2.2e-16). Furthermore, iMARGI is a bulk assay of RNA-chromatin interactions, where the collection of the RNA reads from iMARGI (iMARGI’s RNA reads) measures the transcriptome in the nuclei ^2^. MUSIC’s RNA ensemble also correlated with those of iMARGI’s RNA reads (Figure 2j, rho = 0.9, p-value = 2.2e-16). This indicates that the gene expression levels quantified by MUSIC’s ensemble RNA reads are consistent with those obtained from bulk RNA assays. Moreover, MUSIC detects various types of RNA species (Extended Data Fig. 5c,d), and is strand specific (Figure2k, Extended Data Fig. 5e). Additionally, MUSIC recapitulated the known chromatin association patterns of pre-mRNAs and nuclear-speckle associated RNAs (nsaRNA) (Figure 2l, m, Supplemental Note 6).

## A MUSIC map of human frontal cortex

We generated a MUSIC dataset on 14 postmortem samples of the human frontal cortex (FC) from tissue donors aged 59 and above (Supplementary Table 4). This dataset, hereafter referred to as MUSIC FC, resolved 9,087 single nuclei, 755,123,054 UMNDBC DNA reads and 29,319,780 UMNDBC RNA reads (hg38) (Extended Data Fig. 6a, e). MUSIC FC resolved comparable numbers of single-nuclear RNA reads and DNA-DNA contacts with other methods (Supplemental Note 7, Extended Data Fig. 6b-d).

A clustering analysis based on MUSIC’s single-nucleus RNA reads identified seven cell types (Figure 3a, Supplemental Note 8). The microglia cluster consists of two subclusters, marked by low and high expression levels of Membrane-spanning 4A (MS4A) genes (Extended Data Fig. 7m-o), which may reflect the microglial sub-populations at the chemokine state to the interferon state ^36^. Additionally, a joint analysis of MUSIC FC with a single-nucleus RNA-seq (snRNA-seq) dataset of human frontal cortex ^37^ revealed highly consistent clustering structures and clustering-based cell type assignments between the two datasets (Supplemental Note 9).

**Figure 3.**
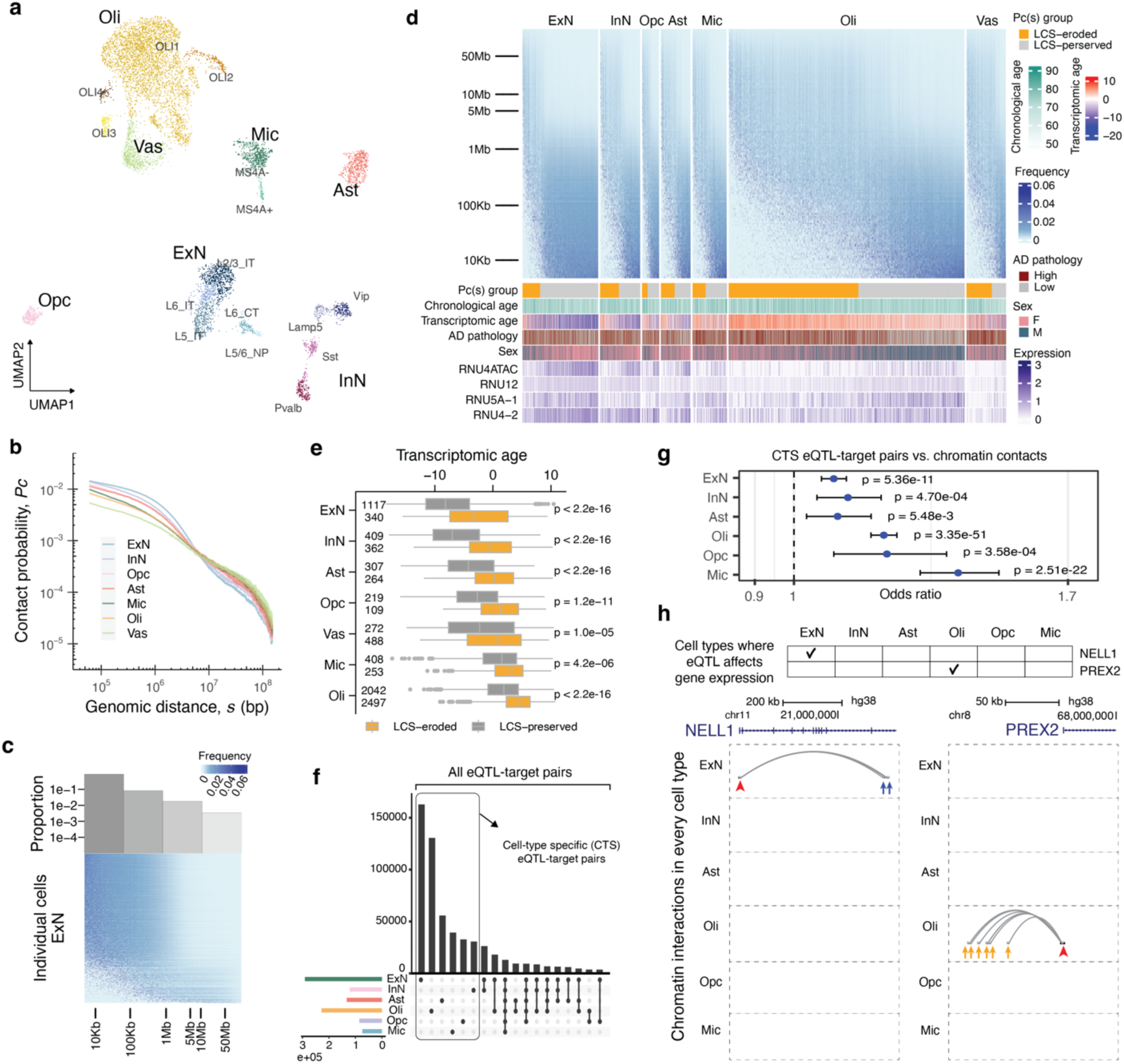
A single-cell map of transcriptome and chromatin conformation in the human frontal cortex. (a) UMAP representation of individual cortical cells based on MUSIC RNA reads. (b) Chromatin contact frequency (Pc, y axis) versus genomic distance (s, x axis) for each cell type (color). (c) Histogram of the proportions of single excitatory neurons (ExN) (rows) with their most frequent chromatin interactions in each genomic bin (x axis) is aligned with the normalized contact frequency (color intensity) vs. genomic distance (x axis) plot for every ExN (row). (d) Chromatin contact frequency vs. genomic distance (y axis) in individual cortical cells (columns). Rows: genomic bins with an exponential size increase. Color scale: chromatin contacts frequency normalized by bin size. Bottom tracks showing the Pc(s) group, Chronological age, Transcriptomic age, AD pathology status, Sex, and the expression levels of several genes. (e) Single cell transcriptomic age (x axis) in each cell type (y axis) colored by chromatin conformation age. Wilcoxon test, one-sided, p-value on the right. Left numbers: sample size. Boxplot: left, center and right edges of box represent the 25th, 50th and 75th quantile. Whiskers extend to 1.5 times the Interquartile Range from the box edges. Data points beyond the whiskers are outliers. (f) Upset plot of the eQTL-target pairs in every cell type. Box indicates the cell-type specific eQTL-target pairs. (g) Association tests between the DD contacts and CTS eQTL-target pairs in every cell type. Center dot: odds ratio. Whiskers: 95% confidence interval of the odds ratio. One-sided Chi-square test, N = 101,785 DD contacts. (h) Examples of CTS eQTL-target pairs supported by cell-type specific DD contacts. Top panel: check mark indicates the cell type (column) where the cis eQTLs of a gene affect this gene’s expression. Bottom panel: genome track view of supporting DDs in every cell type (row), which are the DD contacts (curves) overlapping with the cis eQTLs (blue arrows) and the target gene’s promoter (red arrow).

The stratification analysis by sex (Extended Data Fig. 7a, e) and individual cortex sample (Extended Data Fig. 7d) did not significantly impact the proportions of cells in the clusters or subclusters, except for a higher number of oligodendrocytes in males compared to females (Extended Data Fig. 7b, c, e). Our data indicate a sex difference in the number of cortical oligodendrocytes in the elderly people (>=59 years of age), which aligns with previous studies showing “the lifespan of oligodendrocytes is shorter in females than in males” in mice ^38^. In summary, MUSIC FC data formed clear clusters and these clusters correspond with known cortical cell types and cellular states.

## Heterogeneity in chromatin interactions

Bulk analyses of chromatin conformation revealed that chromatin interaction frequency (Pc) decreases as the genomic distance (s) increases, forming an approximately linear relationship on the log-log scale ^33^. This trend is observed in MUSIC FC as well, where the aggregate chromatin interaction frequency (Pc) in the ensemble of single cells decreases with increasing genomic distance (s) (Figure 3b). Hereafter, we will refer to this trend as the “aggregate Pc-s relationship”.

At the single-cell level, most single cells exhibited a reverse correlation between Pc and s as well, whereas a minority of single cells exhibited the largest Pc not necessarily at the smallest s, a deviation from the aggregate Pc-s relationship (Figure 3c, d). This observation is reminiscent of the recently reported increase in ultra-long-range intrachromosomal interactions during aging in cerebellar granule cells ^39^ (Extended Data Fig. 8e). To test if this observed cellular heterogeneity is compatible with the aggregate Pc-s relationship, we binned the genomic distances and counted the proportion of single cells that exhibit the largest Pc in each genomic distance bin (Figure 3d). The proportion of single cells is smaller in the bins of longer genomic distances, conforming to a reverse correlation that is approximately linear in the log-log scale (Figure 3c). Thus, despite the high degree of cellular heterogeneity, the population summary of the single cells reproduces the previously reported aggregate relationship.

While the different cell types exhibited similar aggregate Pc(s) curves, these Pc(s) curves are not identical (Figure 3b). These differences indicate cell-type variations in chromatin conformation. In particular, excitatory neurons exhibited more frequent chromatin interactions within the sub-Mb range of genomic distances than other cell types (Figure 3b). Consistent with this aggregate behavior, excitatory neurons had a larger proportion of single cells exhibiting the most frequent chromatin interactions at the sub-Mb range compared to the other cell types (Figure 3c, Extended Data Fig. 7f). Together, these observations highlight the influence of cellular composition in each cell type on the cell-type variation in chromatin conformation.

We compared the cellular heterogeneities in Pc(s) and transcriptomic profile. For convenience, we will call the cells with large Pc at small s as ’Local chromatin structure preserved’ (LCS-preserved) cells and those with large Pc at large s as “Local chromatin structure eroded” (LCS-eroded) cells. We simplified the Pc(s) for each cell into a singular score, that is the peak genomic distance of Pc(s), which we call “LCS-erosion score” for this cell. A larger LCS-erosion score reflects a greater loss in the local chromatin structure. In every cell type, we identified the genes with their single-cell expression levels correlated with the single-cell LCS-erosion scores. Pathway enrichment analysis revealed that these genes are enriched in the expected functions of the corresponding cell type (Extended Data Fig. 8a). For example, in excitatory neurons, LCS-eroded cells exhibited reduced expressions of genes associated with Axon guidance, ErbB signaling, Glutamatergic synapse; whereas in inhibitory neurons, LCS-eroded cells exhibited reduced expression of genes in GABA synthesis. These data suggest that the cell-type specific functions are impaired in the LCS-eroded cells.

Next, we analyzed all the cortical cells together, and identified the genes with their single-cell expression levels correlated with the single-cell LCS-erosion scores. A group of small nuclear RNA (snRNA) genes (RNU4ATAC, RNU4-2, RNU5A-1, RNU12) emerged as the group highly correlated with LCS-erosion scores, with high expression in LCS-preserved cells in every cell type (Figure 3d). snRNAs are integral components of the spliceosome, and reduced spliceosome fidelity has emerged as a characteristic of cellular senescence and aging ^40,41^. These data and the cell-type specific analyses let us speculate that the single-cell’s Pc(s) indicates the cell’s “age”. To test this idea, we used the recently published SCALE model ^42^ to compute each single cell’s transcriptomic age, which is a model-based weighted average of the age-related genes’ expression in this cell. LCS-eroded cells exhibited higher transcriptomic ages than the LCS-preserved cells in every cell type, suggesting LCS-eroded cells are older in transcriptomic ages (Figure 3e). In comparison, the chronological age of a sample (age at death) only exhibited weak correlations with LCS-erosion scores (Extended Data Fig. 8b), suggesting the limited ability of the donor’s chronological age to explain the cellular heterogeneity of the LCS-erosion scores (or Pc(s)). Taken together, the cells with reduced chromatin contact frequencies at small genomic distances tend to exhibit older transcriptomic ages.

Seven of the 14 cortex tissues exhibited high pathology of Alzheimer’s disease (AD) (Braak score >= 4), whereas the other seven exhibited low pathology (Braak score <=3). The single cells of the low pathology samples exhibited smaller LCS-erosion scores than those of the high pathology samples in excitatory neurons, inhibitory neurons, astrocytes, oligodendrocytes, and microglia (Extended Data Fig. 8c), revealing a correlation between the loss of local chromatin structure and high pathology. These data are reminiscent of the recent report on the association of a global epigenome dysregulation and AD ^43^. Additionally, a regression analysis reveals the factors with the largest correlations to LCS erosion score include transcriptomic age, cell type, AD pathology, and sex (Supplemental Note 10).

## Correlation of eQTL and chromatin contacts

Cis expression quantitative trait loci (cis eQTLs) represent genomic regions where individual sequence variations contribute to the expression variability of nearby genes ^44^. A recent study reported 481,888 cis eQTLs in the human brain ^44^. Notably, the majority of these cis eQTLs exert their influence on the expression of specific target genes within particular cell types, indicating a strong cell-type specificity in eQTL-target pairings (Figure 3f). The underlying factor responsible for the cell-type specificity observed in eQTL-target pairings remains elusive. We compared the cell-type variations in chromatin contacts (Figure 3b, Supplementary Fig. 3a) with eQTL-target pairings in an association test. This analysis involved all eQTL-target pairs and all the chromatin contacts overlapping with an eQTL and its target gene promoter (supporting DDs), irrespective of their appearance in the same cell type. This test identified a significant enrichment of supporting DDs from the same cell type as the eQTL-target pair (p-value < 2.2e-16, Chi-square test, degrees of freedom = 25) (Supplementary Fig. 3b).

Further analysis focusing on eQTL-target pairs exclusive to individual cell types (cell type-specific (CTS) eQTL-target pairs) reiterated this observation, demonstrating a similar enrichment of supporting DDs within corresponding cell types (p-value < 2.2e-16, Chi-square test, degrees of freedom = 25) (Figure 3g). For example, DD contacts linking NELL1’s cis eQTLs to its promoter were exclusively observed in excitatory neurons, where these eQTLs singularly influence NELL1 expression variability, while similar cell-specific connections were identified for PREX2 in oligodendrocytes, where the eQTLs exclusively impact PREX2 expression variability (Figure 3h). Taken together, the cell types exhibiting chromatin contact between a cis eQTL and its target gene promoter tend to coincide with those where the cis eQTL influences the expression of the target gene.

## Cellular variation in XIST-chrX contacts

The XIST lncRNA is detected in female cortical cells but not in any male cells (Figure 4a), which is consistent with its expected presence in female somatic tissues and absence in male tissues ^45^. In the ensemble of female cells, the XIST lncRNA exhibited a strong association with the entire X chromosome (Figure 4b, c), consistent with its known ability to spread across one of the X chromosomes (the Xi chromosome) ^46,47^.

**Figure 4.**
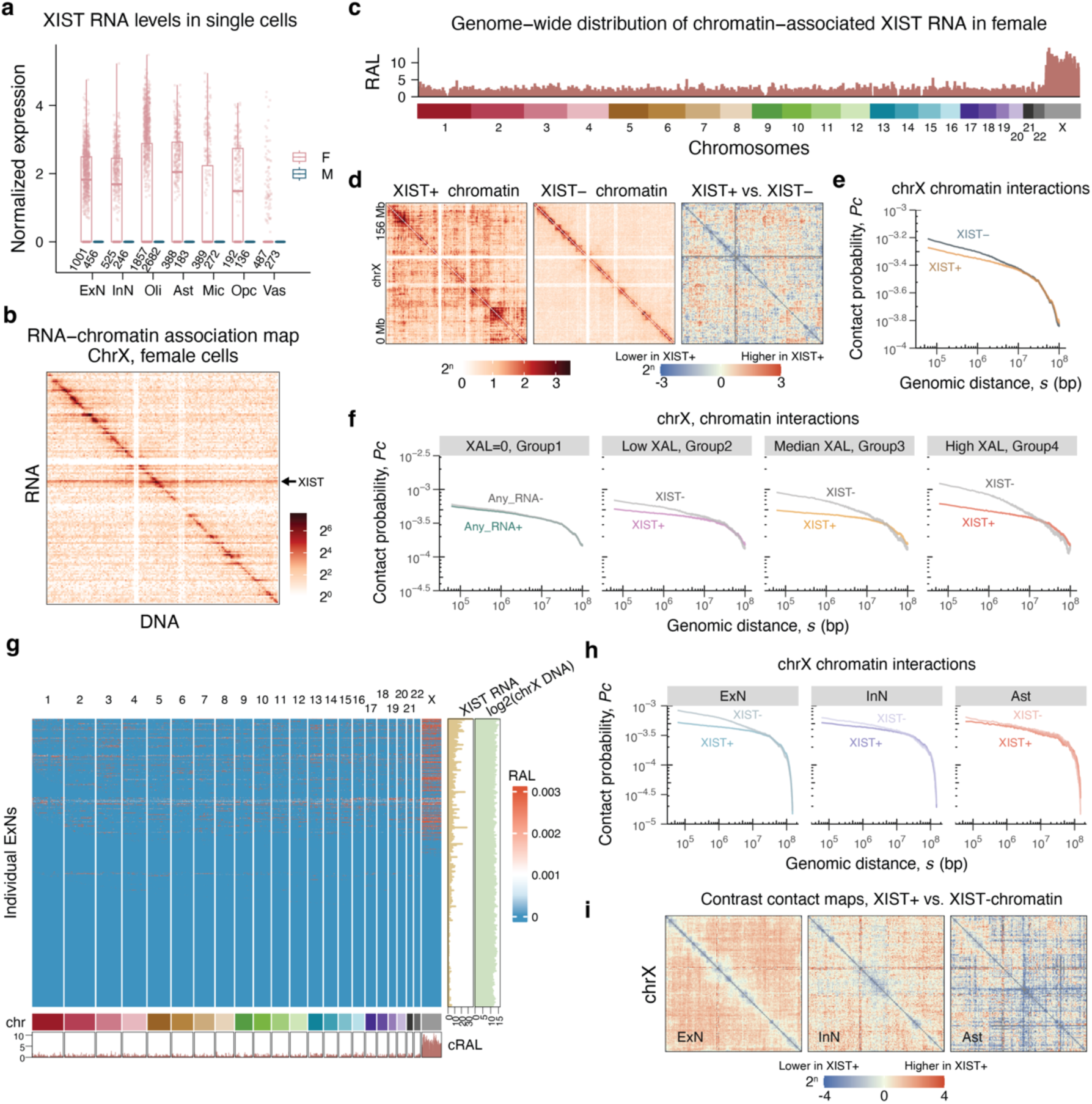
Cellular heterogeneity of XIST-chromatin interactions in female frontal cortex. (a) XIST expression level (y axis) of every single cell (dot) in each cell type for the female and male. Boxplot: bottom, center and top edges of box represent the 25th, 50th and 75th quantile. Whiskers extend to 1.5 times the Interquartile Range from the box edges. Data points beyond the whiskers are outliers. (b) RNA-DNA contact map for Chromosome X based on the ensemble female cells. Each pixel represents the amount of RNA that is transcribed from the genomic bin (row) and is associated with the genomic bin (column). Resolution: 1Mb. (c) Distribution of chromatin-associated XIST on every chromosome in the ensemble of female cells. Resolution: 1Mb. Y axis: the RNA attachment level (RAL) in a genomic bin among the female cells. (d) X chromosome contact maps for XIST-associated (XIST+), not-associated (XIST-) chromatin in female cells, and their contrast. (e) Pc(s) curves showing the frequency of chromatin contacts (Pc, y axis) vs. genomic distance (s, x axis) for XIST+ and XIST-female chrX chromatin complexes. (f) ChrX Pc(s) curves in four female cell groups, with zero, low, medium, and high XAL (left to right). The Pc(s) curves for chromatin with (Any_RNA+) and without any associated RNA (Any_RNA-) do not exhibit a notable difference. The difference between XIST+ and XIST-Pc(s) curves increases as XAL increases. (g) Genome-wide distribution of XIST-chromatin association in ExNs. Each row corresponds to a single ExN, displaying XIST lncRNA attachment levels (red color intensity) across genomic regions on various chromosomes. The bottom track illustrates cumulative XIST RNA attachment levels. Right tracks show XIST RNA read count and X chromosomal DNA read counts in log2 scale. (h) X chromosome Pc(s) curves in female excitatory neurons (ExN), inhibitory neurons (InN), and astrocytes (Ast). The difference between XIST+ (darker color) and XIST-Pc(s) (lighter color) curves is most pronounced in ExN. (i) Contrast contact maps between XIST+ and XIST-chromatins on chromosome X in female ExN, InN, and Ast.

At the single-cell level, the female cortical cells exhibited heterogeneous <u>X</u>IST <u>a</u>ssociation level (XAL), as measured by the number of RD clusters involving XIST lncRNA and X chromosomal DNA in a nucleus (Extended Data Fig. 9c). Recognizing the potential false negatives, we exercised caution in data analysis and interpretation (Supplemental Note 11, Extended Data Fig. 9a). Filtering the female cells based on a threshold of total RNA reads per cell (> 5000) did not eliminate cellular heterogeneity, indicating that the observed heterogeneity cannot be solely attributed to the limited sensitivity of the technique. As expected, the XIST RNA read count in a cell correlated with XAL among the female cells (XIST_ct column vs. Heatmap, Extended Data Fig. 9e), whereas the number of chromosome X DNA reads remained relatively invariant, confirming that the total DNA read count of the X chromosome is independent of XAL (log2 chrX DNA column, Extended Data Fig. 9e). Furthermore, the loss of XIST-chrX association in single female cells correlates with larger sex-difference in the gene expression in the human frontal cortex (Supplemental Note 12).

We compared the X-chromosomal clusters associated with XIST lncRNA (XIST+) and those not associated with XIST lncRNA (XIST-). XIST+ clusters included RD clusters with at least one XIST RNA read, while XIST-clusters were RD and DD clusters that did not contain any XIST RNA read. At the chromosomal scale, most DNA-DNA contacts in XIST-clusters were concentrated near the diagonal line in the chromatin contact map (Figure 4d), similar to the contact maps of autosomes. However, XIST+ clusters exhibited not only near-diagonal contacts but also a significant number of contacts spanning distances of 10 Mb or more (Figure 4d).

Consistent with the chromatin contact maps, the frequency of chromatin contacts (Pc) of XIST-clusters is greater than that of the XIST+ clusters when the genomic distance (s) is smaller than ∼10Mb (Figure 4e). To determine if this separation in Pc(s) curves could be attributed to limited sensitivity in detecting XAL at the single-cell level, a stratification analysis was performed. Female cells were stratified into four groups based on zero, low, medium, and high XALs. Of note, the zero XAL group (Group 1) only contains XIST-clusters, and Groups 2, 3, and 4 contain the same number of cells. Pc(s) of the XIST-clusters (XIST-Pc(s)) is above XIST+ Pc(s) when s is smaller than ∼10Mb in Groups 2, 3, and 4 (Figure 4f). Importantly, the difference between XIST-and XIST+ Pc(s) curves increased from Group 2 to Group 4, indicating that higher XALs in a cell group led to a more pronounced chromatin conformation difference between XIST-and XIST+ clusters. As a control, the Pc(s) curves of X chromosomal clusters associated with or without any RNA (Any_RNA+ Pc(s) or Any_RNA-Pc(s)) in Group 1 (zero XAL group) were nearly indistinguishable (Figure 4f). These data suggest that the active X chromosome (Xa) has a higher contact frequency than the inactive X chromosome (Xi) in the sub-10 Mb range of genomic distances in the female human cortex.

Different cell types exhibited different proportions of XAL-positive (XAL+) cells (Extended Data Fig. 9b), with excitatory neurons having the highest proportion of XAL+ cells (Chi-square, p-value < 10e-16) (Figure 4g). Considering the larger separation between XIST+ and XIST-Pc(s) curves (ΔPc(s)) in XAL-high cells, we anticipated seeing a cell-type difference in ΔPc(s), particularly with excitatory neurons to exhibit a larger ΔPc(s) compared to other cell types. To test this idea, we compared the three cell types with at least 45% of cells exhibiting non-zero XAL, namely excitatory neurons (ExN), inhibitory neurons (InN), and astrocytes (Ast) (Extended Data Fig. 9b). As expected, ExN displayed a larger ΔPc(s) and more long-range interactions than InN and Ast (Figure 4h, i). Specifically, ExN’s XIST+ Ps(c) curve is beneath the XIST-Ps(c) curve when the genomic distance is less than ∼10Mb and traverses above the XIST-Ps(c) curve at ∼10Mb. Although this transversion was consistently observed in InN and Ast, the gaps between the Ps(c) curves were narrower in InN and Ast. Downsampling ExN, InN, and Ast to the same number of cells did not change this observation (Extended Data Fig. 9g). Consistently, seqFISH+ analysis manifested a larger conformational difference between Xa and Xi in ExN compared to InN and Ast in female mice ^48^ (Supplemental Note 13). Taken together, these data suggest a conserved cell-type variation in the spatial organization of the two X chromosomes in the female cortex in mice and humans.

## Discussion

Human cortical cells exhibited a heterogeneity in the distribution of genomic-distance-dependent chromatin contact frequency (Pc(s)). To capture this diversity, we introduced a metric called the LCS-erosion score, designed to condense Pc(s) into a singular value. A higher LCS-erosion score signifies a more pronounced decline in local chromatin contacts. Notably, cells with elevated LCS-erosion scores (termed LCS-erosion cells) demonstrated a tendency toward exhibiting transcriptomic profiles indicative of an ’older’ cellular age in comparison to their counterparts, termed LCS-preserved cells. This distinction sheds light on the correlation between a cell’s chromatin conformation and its transcriptomic age, thus extending our understanding from prior observations associating chromatin structural decline with aging ^49^ to the single-cell level. Consequently, we introduce the concept of a cell’s ’chromatin conformational age’, indicated by the LCS-erosion score.

Compared to the males, female cortexes exhibit less “old” neurons and more “old” oligodendrocytes in chromatin conformation age (Figure 3d, Extended Data Fig. 8d), i.e., a larger oligodendrocyte/neuron ratio in the “old” cells (odds ratio = 6.85, p-value < 2.2e-16, Chi-square test). Reminiscent to this observation, compared to males, female mice exhibit more age-related cell deaths in oligodendrocytes but not in neurons ^38^. Healthy oligodendrocytes protect the normal neuronal activities, and a balance between neurons and oligodendrocytes is required for maintaining their bidirectional communications which ensure the necessary protectivity ^50,51^. We speculate that the disproportionately more “old” oligodendrocytes in females contribute to explain the increased risks in neurodegenerative and mental disorders in women.

The genomic sequence at each expression quantitative trait locus (eQTL) usually remains constant across various cell types within an individual, raising questions about why most eQTLs exert their influence on gene expression in specific cell types. Our findings reveal a connection between cell-type-specific pairing of eQTLs with their target genes and the variability in chromatin contacts between the eQTL and the target gene’s promoter. This underscores the significance of cell-type differences in chromatin contacts for future investigations aimed at unraveling this puzzle.

The widespread impact of 3D genome organization on the expression of numerous genes (Model 1) and the reciprocal influence of gene expression on chromatin conformation (Model 2) continue to be subjects of debate ^2,11^. Recent findings suggest that alterations in chromatin conformation precede changes in gene expression during development, lending support to Model 1 while challenging Model 2 ^19^. However, experiments involving the degradation of cohesin, a pivotal regulator of chromatin conformation, demonstrated limited impact on the expression levels of most genes in a human cell line, casting doubt on Model 1 ^11^. Our data potentially reconcile the limited impact of cohesin degradation with Model 1. The correlation observed between variations in cell-type-specific chromatin contacts and eQTL-target pairing highlights the potential significance of incorporating cell-type and interindividual variations to comprehend a more comprehensive impact.

Following initial debates ^52^, caRNA has been increasingly acknowledged as a structural component of chromatin ^23^. Work in *Drosophila* and *Gallus gallus* suggested 2%–5% of total chromatin-associated nucleic acids are RNA ^53^. In the human MUSIC data, RNA reads account for approximately 4.6% of all the chromatin-derived reads, including all the DD and RD clusters, indicating a relatively consistent proportion of RNA in chromatin-associated nucleic acids across species. Interestingly, approximately 11.7% of the non-singleton chromatin clusters (DD or RD) contain RNA reads. The chromatin clusters containing RNA more frequently demonstrate multiplex DNA-DNA contacts than those devoid of RNA, as shown in Extended Data Fig. 4c and d. This observation aligns with the hypothesis that RNA contributes to spatial genome compartmentalization ^1,2^.

Women exhibit numerous differences in neurodegenerative diseases and mental disorders from men, including that there are twice as many women with late-onset Alzheimer’s disease (LOAD) as men, and that women have a significantly higher frequency of adulthood depression and anxiety compared to men. Notably, many X-linked genes are expressed in the brain and have a role in cognitive functions ^54^. MUSIC data revealed that in female cortical cells, the diminishing association between XIST and chromosome X correlates with reduced structural differences between the active and inactive X chromosomes, and this is associated with greater differences in chrX gene expression between sexes. These multimodal single-cell data provide a critical resource for future investigations on sex differences in health and disease. In summary, MUSIC provides a unique tool to jointly analyze gene expression, multiplex chromatin interactions, and RNA-chromatin associations with single-cell resolution from complex tissue.

## Supporting information

Full supplementary file

## METHODS

### Critical Reagents

#### The RNA Linker

The RNA Linker is a single stranded chimeric oligonucleotide with 17 DNA nucleotides at its 5’ end (ssDNA, Extended Data Fig. 2e) and 10 RNA nucleotides at its 3’ end (ssRNA, Extended Data Fig. 2e), denoted as 5’ /5OH/CGAGGAGCGCTTNNNNNrArUrArGrCrArUrUrGrC/3OH/ 3’, where A, C, G, T, and N denote DNA nucleotides, rA, rC, rG, rT denote RNA nucleotides, NNNNN denotes 5 randomized DNA nucleotides that serves as a unique molecular identifier (UMI). The RNA Linker was synthesized by IDT.

The RNA Linker is designed for (1) efficient ligation with RNA through the RNA Linker’s ssRNA, (2) efficient ligation with the 1st set of Cell Barcodes through the DNA Linker’s ssDNA, which complements with the 7 nt overhang in the 1st set of Cell Barcodes.

#### The DNA Linker

The DNA Linker is a hybridized product of two DNA strands with the top strand being 5’ /5Phos/CTAGACACTGTGCGTATCTNBAAAAAAAAAAAAAAAAAAAAAAAAAAAAAA/3OH/ 3’, where N denotes a random base and B denotes any base except A, and the bottom strand being 5’ /5OH/CGAGGAGNNNNNACAACGCACAGTGTCTAGT/3OH/ 3’, where NNNNN denotes 5 randomized DNA bases that serve as a UMI. After hybridization, the DNA Linker contains 15bp double-stranded DNA (referred to as dsDNA) and 36 nt (top-ssDNA) and 15 nt unhybridized single stranded DNA (bottom-ssDNA), and a single base (T) overhang at the bottom strand (Extended Data Fig. 2g). The 36 nt top-ssDNA is reverse complementary to the 10X Barcodes in the Chromium Next GEM Single Cell 3’ Reagent kit (PN-1000268). The two strands of the DNA Linker are synthesized by IDT.

The DNA Linker is designed for (1) efficient ligation with fragmented chromosomal DNA through the DNA Linker’s 1 nt overhang by a sticky-end ligation, (2) efficient ligation with the Cell Barcodes, through the DNA Linker’s 15 nt bottom-ssDNA, and (3) efficient hybridation with the 10X Barcodes through the polyA sequence within the top-ssDNA.

#### The Cell Barcodes

The Cell Barcodes contain three sets of barcodes, referred to as the 1st, 2nd, and 3rd sets of Cell Barcodes. Every set of Cell Barcodes has three components, namely a 7 nt top-strand overhang, a 14 bp dsDNA region, and a bottom-strand overhang (7 nt for the 1st and 2nd sets, 11 nt for the 3rd set). The 14 bp dsDNA region contains a unique sequence to every Cell Barcode (double-stranded N14, Extended Data Fig. 2d, i). Every set of Cell Barcodes contains 96 unique barcodes, where each barcode is unique in this 14 bp dsDNA (Supplementary Table 2).

In the current version of MUSIC (v1.0), each set of Cell Barcodes contains 96 unique barcodes based on their dsDNA regions, resulting in a total of 884,736 unique sequence combinations. The three sets of Cell Barcodes are designed to maximize the ligation efficiency for sequentially ligating the 1st set of Cell Barcodes with the RNA Linker and the DNA Linker, the 2nd set with the 1st set, and the 3rd set with the 2nd set of Cell Barcodes. The optimal ligation efficiency is achieved by the complementarity of the overhang sequences (Extended Data Fig. 2d, i). Out-of-order ligations such as a ligation of the 3rd set with the 1st set of Cell Barcodes are minimized because the overhang of the 3rd set does not complement with the overhang of the 1st set of Cell Barcodes.

Additionally, the 3rd set of Cell Barcodes is also designed to complement the 22 nt sequence at the 3’ side of the Index Adaptors.

The 1st and 2nd sets of Cell Barcodes are synthesized by Sigma-Aldrich. The 3rd set of Cell Barcodes are synthesized by IDT.

#### 10X Barcodes

The 10X Barcodes are included in the Chromium Next GEM Single Cell 3’ Reagent kit PN-1000268 from 10X Genomics. Each 10X Barcode is an 82 nt oligonucleotide with a partial (22 nt) Illumina Read 1 sequence (Read 1), a 16 nt unique barcode sequence (N16, 10X GEM Barcode), a 12 nt unique molecular identifier (N12, 10X UMI), a 30 nt polyT sequence, a V (A or C or G) and an N (any base) (10X Barcode, Extended Data Fig. 2j, k). The 10X GEM Barcode is shared among the barcodes of the same GEM. The 10X UMI is unique to every 10X Barcode.

#### The Index Adaptors

The Index Adaptors contain three segments, namely the 24 nt Illumina P7 sequence, a 8 nt unique identifier sequence called I7, and the 34 nt Illumina Read2 sequence (Index Adaptor, Supplementary Table 2). In this release, MUSIC v1.0 uses eight distinct I7 Barcodes, providing for a total of approximately 83.5 million Complex Barcodes (8 (I7 Barcodes) × ∼3.5 million (10X Barcodes)). Eight Index Adaptors are used for every library construction. Each of the eight Index Adaptors has a unique I7 sequence. We call these eight Index Adaptors a set of Index Adaptors. Each Index Adaptor can hybridize with the complementary Read2 sequence in the 3rd set of Cell Barcodes to start a PCR reaction.

Meanwhile, they serve as sample barcodes. We designed a total of three sets of Index Adaptors to allow for constructing three libraries from three input samples and sequencing them together (Supplementary Table 2). These three sets of Index Adaptors share P7 and Read2 sequences and differ by their I7 sequences. These three sets of Index Adaptors serve as a sample index to differentiate three samples. The Index Adaptors are synthesized by IDT.

#### The Universal Adaptor

The Universal Adaptor contains an Illumina P5 sequence and an Illumina Read1 sequence (Universal Adaptor, Supplementary Table 2). The Universal Adaptor can hybridize with the 10X Barcodes through their complementary Read1 sequence to start a PCR reaction. The Universal Adaptor is synthesized by IDT.

### Cell Culture

H1 human embryonic stem cells and E14 mouse embryonic stem cells were obtained from the 4D Nucleome Consortium and cultured according to the 4D Nucleome Consortium approved protocols (https://www.4dnucleome.org/). Briefly H1 cells were grown at 37°C under 5% CO_2_ on matrigel (Corning, 354277) coated dishes. Cells were maintained in complete mTeSR medium prepared from basal medium (Corning, 85851) with 5X supplement (Corning, 85852). Medium was replaced daily. Cell passage number was kept under P10. E14 cells were cultured on plates coated with 0.1% gelatin (EMD, SF008) in serum free 2i/LIF medium. The serum free 2i/LIF medium was made from the base medium (1:1 mixture of NeuroBasal medium (Gibco, 21103-049) and DMEM/F12 medium (Gibco, 11320-033) supplemented with 0.5X N2 supplement (Gibco, 17502-048), 0.5X B27 supplement (Gibco, 17504-044), and 0.05% BSA fraction V (Gibco, 15260-037)) supplemented with 1 µM PD0325901 (Reprocel,l 04-0006-02C), 3 µM CHIR99021 (Reprocell, 04-0004-02C), 0.15 mM Monothioglycerol (Sigma, M6145-25ML), 1000 U/mL LIF (Cell Guidance Systems, GFM200). The medium was replaced daily. Cell passage number was kept under P10.

### Crosslinking and nuclei isolation for cell lines

After cells become confluent in a 10 cm dish, the media was removed and washed once with PBS. 1 mL of Accutase (EMD, SF006) was added and incubated for 3 min at 37°C to dissociate the cells. 10 mL of PBS was used to generate single cell suspension by pipetting. Cell pellets were formed by centrifugation at 330 x g for 3 min. 10 mL of 2 mM disuccinimidyl glutarate (DSG) dissolved in 1X PBS was added to crosslink and resuspend the cells in a LoBind tube and incubate at room temperature for 45 min with gentle rotation. After incubation, cells were collected by centrifugation at 1,000 x g for 4 min to remove DSG solution and washed once with 1X PBS then centrifuged again at 1,000 x g for 4 min to remove the supernatant. After washing, cells were thoroughly resuspended in 10 mL of 1X PBS containing 3% formaldehyde and incubated for 10 min with gentle rotation. The crosslinking reaction was stopped by adding 3 mL of 2.5 M glycine per 10 mL of 3% formaldehyde and incubating for 5 min with rotation. Cells were then centrifuged at 1,000 x g for 4 min to remove the supernatant. Next, cells were washed twice with ice-cold 1X PBS containing 0.5% BSA (wt/vol) and centrifuged at 1,000 x g for 4 min. After the wash, cells were resuspended in 1X PBS with 0.5% BSA (wt/vol), and each cell aliquot contained 5 million cells in a 1.5 mL tube. The cell pellets were obtained by centrifugation at 1,000 x g for 5 min, snap-frozen in liquid nitrogen, and stored at −80°C.

Frozen cells were thawed on ice and resuspended in 1.4 mL of cell lysis buffer A for every 5 million cells as previously described ^12^ (50 mM HEPES pH 7.4, 1 mM EDTA pH 8.0, 1 mM EGTA pH 8.0, 140 mM NaCl, 0.25% Triton X-100, 0.5% IGEPAL CA-630, 10% glycerol, 1X proteinase inhibitor cocktail (PIC)). For the mixed species experiment, equal amounts of human H1 (2.5 million) and mouse E14 (2.5 million) cells were resuspended together. After a 10-min incubation on ice, cell pellets were collected by centrifugation at 900 x g for 4 min at 4°C. Cell pellets were resuspended in 1.4 mL of cell lysis buffer B (10 mM Tris-HCl pH 8, 1.5 mM EDTA, 1.5 mM EGTA, 200 mM NaCl, 1X PIC) and kept on ice for 10 min. The isolated nuclei were collected at 900 x g for 5 min at 4°C. 200 µL of 1X rCutSmart buffer (NEB, B7204S) containing 0.25% SDS was used to thoroughly resuspend and permeabilize the nuclei and incubated at 62°C for 10 min using Eppendorf Thermomixer C (Eppendorf). After the incubation, 60 µL of 1X rCutSmart buffer containing 10% Triton X-100 (wt/vol) was mixed with the SDS solution, and the reaction was incubated at 37°C for 15 min while shaking at 800 rpm. The treated nuclei were centrifuged at 900 x g for 2 min at 4°C to remove supernatant and washed once with 1X rCutSmart buffer.

### Crosslinking and nuclei isolation for post-mortem brain

Each 50 mg of post-mortem human brain frontal cortex sample was kept on ice in a 1.5ml LoBind tube and chopped into smaller pieces by pestle. The brains samples were transferred into a 15 mL LoBind tube and incubated at room temperature for 45 min with a gentle rotation in 10 mL of 2 mM DSG that was dissolved in 1X PBS. After incubation, the tissue sample was centrifuged at 1,000 x g for 4 min and washed once with 1X PBS to remove DSG solution. After washing, the tissue sample was thoroughly resuspended in 10 mL of 1X PBS containing 3% formaldehyde and incubated for 10 min with a gentle rotation. The crosslinking reaction was stopped by the addition of 4 mL of 1.25 M glycine followed by an incubation of 5 min with a rotation. The tissue sample was then centrifuged at 1,000 x g for 4 min and washed twice with ice-cold 1X PBS containing 0.3% BSA (wt/vol).

Chromium Nuclei Isolation kit (10X genomics, 1000494) was used to isolate nuclei from crosslinked cortex samples according to the section for “single cell gene expression & chromium fixed RNA profiling” (page 25 to page 30 from sample user guide). Specifically, 50mg frozen tissue was placed into a pre-chilled sample dissociation tube. 400 µL of the provided lysis buffer was added to the tube, and tissues were dissociated until homogeneous with plastic pestle. 600 µL lysis buffer was further added into the tube and mixed 10 times by pipetting. After a 10 min incubation on ice, the solution was evenly loaded into two nuclei isolation columns and centrifuged at 16,000 x g for 20 s at 4°C. The flowthrough in the collection tube which contains nuclei was vortexed 10 s at 3,200 rpm to resuspend nuclei. The collection tube was centrifuged for 3 min at 500 x g at 4°C to pellet nuclei, and the supernatant was removed. The nuclei were resuspended in a 500 µL debris removal buffer provided by the kit by pipetting 15 times. The nuclei were centrifuged at 700 x g for 10 min at 4°C, and the supernatant was removed. The nuclei were resuspended in 1 mL of wash and resuspension buffer twice. The supernatant was removed after centrifugation at 500 x g for 5 min at 4°C, which left a purified pellet of isolated nuclei (Supplementary Fig. 1a, b). All the following steps are the same for either cell line or human cortex samples.

### Ligation of the RNA Linker with RNA

The nuclei were resuspended in 250 µL of 5’ phosphorylation master mix (1X T4 PNK buffer, 500 U/mL of T4PNK, 1 mM ATP, 1 U/µL of RNAse inhibitor (Roche, 3335399001)) and incubated at 37°C while rotating at 800 rpm for 1 hour to phosphorylate the 5’ ends of RNA. Nuclei were washed once with PBS Wash Buffer-1 (1X PBS, 1 mM EDTA, 1 mM EGTA and 0.1% Triton X-100) and twice with PBS Wash Buffer-2 (1X PBS, 0.5% BSA (wt/vol) and 0.1% Triton X-100). The RNA Linker is a single stranded chimeric oligo with DNA’s 5’ hydroxyl group end and RNA’ 3’ hydroxyl group (5’-OH-CGAGGAGCGCTTNNNNNrArUrArGrCrArUrUrGrC-OH-3’). An RNA Ligation Mix is made with 4 µM of the RNA linker, 1X T4 RNA ligation buffer, 400 U/mL of T4 RNA ligase 1, 15% PEG 8000, 1 mM of ATP, and 1 U/µL of RNAse inhibitor. The isolated nuclei were thoroughly mixed with 250 µL of the RNA Ligation Mix to ligate the RNA Linker with nuclear RNA. The mixture was incubated at 25°C for 2 hours then 16°C overnight with an intermittent mixing at 800 rpm (30 seconds on and 270 off). After ligation, the nuclei were washed once with PBS Wash Buffer 1 and three times with PBS Wash Buffer 2.

### Chromatin Digestion

All the washed nuclei were resuspended in a digestion master mix (300 µL of 1X rCutSmart buffer containing 30 µL of 5,000 U/mL of HpyCH4V with 1 U/µL of RNAse inhibitor). The digestion master mix was kept for 3 hours at 37°C while rotating at 800 rpm. The nuclei were collected at 900 x g for 2 min with supernatant removed. The nuclei were further washed once with 900 µL of PBS Wash Buffer 1 (1X PBS, 1 mM EDTA, 1 mM EGTA and 0.1% Triton X-100) and three times with 900 µL of PBS Wash Buffer 2 (1X PBS, 0.3% BSA (wt/vol) and 0.1% Triton X-100).

### Ligation of the DNA Linker with DNA

To create sticky end for DNA linker ligation, the nuclei was suspended in 250 µL of dA-tailing reaction master mix (1X NEBNext dA-tailing reaction buffer, 200 U/mL of Klenow fragment, 1 U/µL of RNAse inhibitor) and incubated at 37 °C while rotating at 800 rpm for 1.5 hours. Next, the nuclei were washed once with PBS Wash Buffer-1 and three times with PBS Wash Buffer-2. The DNA Linker is a hybridized product of two DNA strands with the top strand being 5’-Phos-CTAGACACTGTGCGTATCT NBAAAAAAAAAAAAAAAAAAAAAAAAAAAAAA-OH-3’, and the bottom strand being 5’-OH-CGAGGAGNNNNNACAACGCACAGTGTCTAGT-OH-3’. The DNA Linker contains 14 bp double-stranded DNA (referred to as dsDNA) and 36 nt and 15 nt unhybridized single stranded region which is referred to as the (top-ssDNA and bottom-ssDNA) (Extended Data Fig. 2g, h). A DNA Ligation Master Mix is made of 4.5 µM of the DNA Linker, 0.2X of 2X Instant Sticky-end Ligase Master Mix (NEB, M0370), 0.8X of 5X Quick Ligase Buffer (NEB, B6058S), 6% (vol/vol) of 1,2-propanediol (Sigma-Aldrich, 398039), and 1 U/µL of RNAse inhibitor). To ligate the DNA Linker with the sticky end DNA, all the nuclei were thoroughly mixed with 250 µL of the DNA Ligation Master Mix. The ligation reaction was kept at 20°C for 6 hours with an intermittent mixing at 1600 rpm (30 seconds on and 270 seconds off).

### Ligation of Cell Barcodes

To phosphorylate the 5’ end of the Linker, the nuclei were resuspended in 250 µL of 5’ phosphorylation master mix and incubated at 37°C while rotating at 800 rpm for 1 hour. Afterwards, the nuclei were washed once with PBS Wash Buffer 1 and three timeswith PBS Wash Buffer 2. The nuclei were resuspended in 900 µL of PBS Wash Buffer 2 with 0.2 U/µL of RNase Inhibitor and filtered through a 10 µM cell strainer (pluriStrainer, 43-10010-50). 6 µL of the nuclei suspension was stained with 6 µL of Ethidium homodimer-1, and the number of nuclei was counted by Countess II Automated Cell Counter (ThermoFisher). A total of 288 barcodes were taken from Hawkins et al^2^ and split to three barcode sets, named Barcode Set 1, 2, and 3. Each barcode takes the form of 7 nt_overhang-dsDNA-7 nt_overhang (Extended Data Fig. 2d, i, Supplementary Table 2). These 288 barcodes are collected called the Cell Barcodes. Three Ligation Master Mixes are prepared, with each Ligation Master Mix containing one set of barcodes (5.4 µM) and 0.2X of 2X Instant Sticky-end Ligase Master Mix, 0.8X of 5X Quick Ligase Buffer, 6% (vol/vol) of 1,2-propanediol, 0.8 U/µL of RNAse inhibitor. The three Ligation Master Mixes are named Ligation Master Mix 1, 2, and 3, corresponding to Barcode Set 1, 2, and 3, respectively.

#### The first round of split-pool

Up to 100,000 nuclei were collected for split-pool to ensure the majority of nuclei were labeled with unique cell barcode combinations. The nuclei suspension was filled to 1144 µL with PBS Wash Buffer 2 and 24 µL of RNAse inhibitor and subsequently split into 96 wells. To ligate Barcode Set 1 with the RNA Linker and the DNA linker, the nuclei in each well are incubated in the Ligation Master Mix 1 at 20°C overnight with an intermittent mixing at 1600 rpm (30 seconds on and 270 off). After overnight incubation, the reaction was quenched by an addition of 60 µL of quenching buffer (1X PBS, 50 mM EDTA, 50 mM EGTA, 0.1% Triton X-100) and incubated for 10 min at 20°C. The nuclei solutions from the 96 wells were pooled together into a 15 mL LoBind tube. 95 µL of quenching buffer was added to each well to rinse and collect any remaining nuclei and pooled into the same 15 mL tube. The nuclei were centrifuged at 900 x g for 4 min, and the nuclei were transferred into a 1.5 mL tube with 0.5 mL of remaining supernatant. 500 µL of PBS Wash Buffer 2 was used to rinse the 15 mL tube and collect the residual nuclei into the same 1.5 mL tube. The nuclei were washed three times with 900 µL of PBS wash buffer 2 by the centrifugation at 900 x g for 2 min.

#### The 2^nd^ and 3^rd^ round of split-pool

The pooled nuclei were subjected to the same split-pool procedure as the first round, except that the Ligation Master Mixes 2 and 3 were used for the 2^nd^ and the 3^rd^ rounds, replacing the Ligation Master Mix 1.

#### Addition of Complex Barcodes

We use a combination of two sets of the barcodes to jointly differentiate individual molecular complexes. This combination is hereafter referred to as the Complex Barcodes. The first set of barcodes are those 3.5 million oligos provided in the Chromium Next GEM Single Cell 3’ Reagent kit PN-1000268 (10X Barcodes). Each oligo is an 82-base oligonucleotide with 16 nt barcode and 12 nt unique molecular identifier (10X BC+UMI, Extended Data Fig. 2j, k). The second set of barcodes is composed of 8 barcodes (Index Barcodes). Each barcode is 8 nt (i7 in Extended Data Fig. 1h, Supplementary Table 2).

The nuclei were resuspended in 250 µL of 3’ Dephosphorylation Buffer (1X PNK buffer, 0.5 U/µL T4PNK, 1 U/µL of RNAse inhibitor) and incubated at 37°C for 1 hour with rotating at 800 rpm to convert any 2’, 3’ cyclic phosphate on RNA to 3’-OH. The nuclei were washed once with PBS Wash Buffer 1 and three times with PBS Wash Buffer 2 and centrifuged at 900 x g for 2 min. To add polyA sequences to all the RNA molecules, the nuclei were resuspended in PolyA Tailing Buffer (1X E. coli Poly(A) Polymerase Reaction Buffer, 0.08 U/µL of E. coli Poly(A) Polymerase, 1 mM of ATP, RNAse inhibitor 1U/µL). The mixture was kept at 37°C for 10 min while rotating at 800 rpm. After adding polyA tails, the nuclei were thoroughly resuspended in 1X PBS with 0.04% BSA (wt/vol) and filtered through a 10 µM cell strainer (pluriStrainer, 43-10010-50) into 1.5 mL tube to obtain isolated nuclei. 6 µL of the nuclei-containing solution was stained with 6 µL of Ethidium homodimer-1 and counted by Countess II Automated Cell Counter. 5,000 single nuclei were transferred to a Covaris microtube-15 and filled to 15 µL with 1X PBS with 0.04% BSA (wt/vol). The nuclei were sonicated using Covaris M220 Focused-ultrasonicator with water temperature 6°C, incident power 50 W, duty factor 5 for 5 min to release chromatin complexes.

To add the 10X barcodes to the polyadenylated RNA and the top-ssEND of the DNA Linker, the sonicated nuclei complexes were transferred into a 1.5mL LoBind tube and mixed with 25 µL of water, 18.8 µL of RT reagent B, 2 µL of Reducing Agent B, 8.7 µL of RT enzyme C. The mixture was transferred to one well in ChIP G, which is loaded to the 10X Chromium controller according to Steps 1.1 to 1.5 in the protocol of Chromium Next GEM Single Cell 3’ Reagent kit. The retrieved droplets were transferred into a PCR tube for cDNA synthesis according to the 10X protocol. The droplets were broken, and the aqueous phase was obtained according to Step 2.1 in the 10X protocol.

The aqueous phase containing nucleic acids were filled to 200 µL with nuclease free water and split into eight aliquots in LoBind 1.5 mL tubes. 25 µL of 2X reverse crosslinking buffer (400 mM NaCl, 0.4% SDS, 50 mM EDTA, 50 mM EGTA, 0.04 U/µL Proteinase K) was added into each tube, and reverse crosslinking reaction was incubated at 50°C for 2 h and 55°C overnight with shaking at 800 rpm. In each aliquot, the reverse crosslinked nucleic acids were purified with Monarch RNA purification kit (NEB, 76307-460) and eluted into 21 µL of nuclease free water. The eluted DNA and RNA molecules were incubated at 55°C for 15 min with 1X isothermal amplification buffer II, 0.32 U/µL of Bst 3.0 DNA polymerase, additional 6 mM MgSO4, 1.4 mM dNTP Mix, and 0.5 U/µL of RNasin. The product was purified with 1.8X RNA clean Ampure beads (Beckman Coulter life science, A63881) and eluted into 20 µL of nuclease free water. PCR is carried out for each aliquot with 2.5 µL of 10 µM shared Universal Adaptor (P5 and Read1 in Extended Data Fig. 1h) and 2.5 µL of 10 µM aliquot-specific primer in 25 µL of NEBNext Ultra II Q5 Master Mix. The aliquot-specific primers are the eight Index Adaptors (see Reagents created by this work). PCR is carried out at 13 to 14 cycles. The amplified DNA was purified with 1.2X Ampure beads and eluted into 12.5 µL of nuclease free water. The purified DNA solutions from the eight aliquots were combined and loaded into 5 lanes of 4% E-gel (Invitrogen, G401004). The DNA bands with the size between 300 bp and 1.2 Kb were excised. The DNA was extracted with NEB Monarch gel purification kit (NEB, T1020S) using two columns and eluted in 30 µL of the elution buffer.

### Sequencing

The molarity of the sequencing library was measured by Qubit 4.0 Fluorometer (Invitrogen, Q33238) using Qubit dsDNA HS assay kit (Invitrogen, Q33231). Fragment size distribution was assessed by Agilent bioanalyzer with High sensitivity DNA ChIP. The library was sequenced by UC San Diego IGM Genomics Center utilizing an Illumina NovaSeq 6000. The sequencer is set to read a 28 bp sequence next to the Universal Adapter as the Read1, a 8 bp index sequence from the I7 region inside the Index Adapter, and a 150 bp sequence next to the Index Adapter as the Read2.

### Computational analysis

#### The MUSIC-docker data processing pipeline

We developed MUSIC-docker to process MUSIC sequencing data using Docker to encapsulate a Snakemake ^55^ pipeline, ensuring cross-platform execution. It handles i7 index-splited paired-end fastq files, processes them into RNA and DNA sequences separately, adds cell and complex barcodes, and maps to the genome, removing PCR duplicates and deriving processed files. Detailed documentation is available at: http://sysbiocomp.ucsd.edu/public/wenxingzhao/MUSIC_docker/intro.html, and Supplementary Note 14.

The raw sequencing output (.bcl) is converted to FASTQ files with bcl2fastq, producing eight FASTQ files with 28bp Read1 and 150bp Read2. Read1 includes the 10X Barcode and 10X UMI, while Read2 contains Cell Barcodes, RNA/DNA Linker, and RNA/DNA insert sequences.

Demultiplexing step extracts cell and complex barcodes information and fragment sequence information, creating separate FASTQ files for RNA and DNA reads individually where the read sequence will be the insert, and the read name will be the fragment identity information.

To address potential artificial sequences introduced by sequencing errors and experimental design, we remove consecutive As or Gs from the 3’ of the DNA inserts if their length is longer than 20bp. For RNA reads, we will detect the ssDNA region of the RNA Linker sequence “CGAGGAGCGCTT” and remove any sequence following it. We use cutadaptor (v2.8) ^56^ with parameter ‘-q 15 -m 20‘. DNA and RNA inserts are mapped to the genome using bowtie2 (v5.4.0) ^57^ with parameters “bowtie2 -p 10 -t --phred33 -x ” and bwa (v0.7.17) ^58^ mem with parameters “-SP5M”, respectively. Uniquely mapped reads are selected for downstream analysis.

After reads mapping, PCR duplicates, identified by shared 10X UMI and mapped coordinates are removed using customized script. The script sorts the BAM file, scans it once, and flags duplicates if they meet specific criteria: 1) it maps to a location within 8 base pairs of the previous read, 2) it shares the same cell barcode and molecular barcode with the previous read, and 3) its UMI exhibits a Levenshtein distance of less than 2 base pairs from the UMI of the previous read. All identified PCR duplicates are subsequently removed to ensure the integrity of downstream analyses.

Finally, deduplicated BAM files from each I7 index library are merged into a comprehensive, sorted BAM file, capturing essential information for both DNA and RNA, including cell barcode, molecular barcode (10X and I7 index), and insert mapping location.

#### Mix species experiment

Based on the published method ^12^, we assigned a cell to a species if 95% of all its DNA reads could map to a single species. Cells with uniquely mapped non-duplicated DNA reads less than 1000 are classified as ambient cells. To calculate the single cluster mix species rate, we assigned a cluster to a single species if more than 99% of its uniquely mapped non-duplicated DNA reads come from that species.

#### Parsing multiplex clusters into pairwise interactions with normalization

Each cluster is a collection of reads that share the same Cell Barcodes, 10X Barcode and I7 Barcode (CB+GEM+I7). A cluster of two reads (cluster size = 2) corresponds to a pairwise interaction. A cluster of three or more reads corresponds to a multiplex cluster. Each multiplex cluster is decomposed into pairwise interactions with a normalization procedure that adjusts for the total number of combinations of pairwise interactions from a multiplex cluster as previously described ^12,22^. For homotypical clusters, which are clusters containing exclusively DNA reads or RNA reads, each homological cluster of size N is first decomposed into all non-overlapping pairs, and then each pair is normalized by a factor of 1/N. For heterotypic clusters, which are clusters containing both DNA and RNA reads, each heterotypic cluster is first decomposed into all DNA-RNA pairs, and then each pair is normalized by a factor of 1/(M+N) where M and N are the numbers of DNA and RNA reads. This normalization removes the difference in the number of pairwise decompositions from different-sized clusters, thus ensuring the larger clusters do not get an inflation in the number of decomposed pairwise interactions ^12,22^.

To generate a two-dimensional contact map of DNA-DNA interactions, we first define the size of each unit, typically represented as genomic bins, for rows and columns. The weight assigned to bin [i, j] is determined by the total number of clusters that contain DNA reads mapped to both the ith and jth bins, and it is normalized by the cluster size as previously described. To compare DNA-DNA contact differences within a specific genomic region, cluster sizes are calculated using only DNA reads mapped to that region, considering small, medium, and large clusters. If the DNA-DNA contact map is generated for multiple cells, the weighted sum of all clusters is calculated. Similar methods are applied to derive RNA-DNA 2D contact maps, with the calculation of the weight of each interaction adjusted accordingly.

To determine the RNA attachment level (RAL) at specific genomic bins, representing 1D RNA-chromatin contacts for a particular RNA of interest, we calculate the total weighted RNA-DNA interactions involving the RNA of interest and DNA ends mapped to the desired genomic bins. For ensemble maps, RAL values from individual cells are summed.

Finally, for visualizing the two-dimensional contact maps of DNA-DNA or RNA-DNA interactions, the raw contacts, obtained following the previously mentioned procedures, are scaled up using a linear factor, typically 100 or 1000. This amplification step prevents the application of logarithmic transformation to decimal values. After amplification, a logarithmic transformation is performed to enhance visualization of the contact maps.

#### Comparison between MUSIC DNA-DNA contacts with Micro-C data

In order to achieve consistency in the total contacts scale between MUSIC and Micro-C, contact maps obtained from both methods underwent a standardized transformation. Initially, the contact maps derived from Micro-C or MUSIC were logarithmically scaled. Subsequently, the scaled contact maps were normalized to their respective maximum values, resulting in all contacts being constrained within the range of [0, 1]. The Micro-C data used in this study was obtained from the 4DN portal ^59^ (4DNFI2TK7L2F) ^34^. To extract the raw contacts from the .hic file, the straw ^60^ tool was employed.

To calculate the compartment score (PC1 score) for both Micro-C and MUSIC DNA-DNA contact matrices, we first computed the Expected Contact Matrix. Subsequently, we determined the PC1 score from the correlation matrix of the Observed/Expected ratio matrices, where the Expected Matrix was derived by calculating average contact frequencies as a function of genomic distances. For the comparison of PC1 correlations between Micro-C and MUSIC, the calculations were performed individually on each chromosome. The reported correlation represents the median of these correlations across all chromosomes.

#### nsaRNA and pre-mRNA RAL

Pre-mRNAs are RNA reads that exhibit an overlap of at least 15 base pairs with gene introns and are classified as protein-coding RNAs. The calculation of the RD (RNA-DNA) cluster weight involves the inclusion of all nsaRNAs and pre-mRNAs, along with their associated DNA reads. RD clusters with a cluster size not exceeding 1000 were selected for the computation of the RNA attachment level (RAL). The H1 A/B compartment data used in this study was obtained from the 4DN data portal (4DNFID162B9J) ^2^.

#### Calculate genomic distance versus contact frequency curve from MUSIC data

The relationship between contact frequency and genomic distances within chromosomal arms was systematically examined. Initially, the genomic distance range from 10 base pairs to 150 megabases was divided into 500,000 equally sized bins. Subsequently, for DNA reads originating from DD or RD clusters, the genomic distances of all intrachromosomal pairwise interactions were determined, and the frequencies for each genomic distance bin were computed. These frequencies were then normalized based on the weight assigned to each interaction. The normalized frequencies were calculated for each genomic bin in every cluster and single cell. Finally, the genomic distances versus contact frequencies were aggregated across all clusters from all single cells.

We used cooltools (0.5.4) ^61^ for generating the genomic distance versus contact frequency curve in Micro-C data. We downloaded the .mcool file for H1 Micro-C from the 4DN data portal (4DNFI9GMP2J8) ^34^.

#### Brain single cells preprocessing and filtering

For each brain sample we applied standard MUSIC_tool docker pipelines to get valid RNA and DNA reads information for each single cell. To select high quality single cells for robust analysis and interpretation of our data, we removed cells that have less than 100 RNA reads or less than 5000 DNA reads for downstream analysis on brain samples.

#### Brain sample transcriptome merging

We first constructed the single cell RNA expression count matrix by calculating the number of RNA reads mapped to each human gene (GENCODE ^62^, v36, chrM genes are excluded) for all single cells from all 14 brain samples. We then constructed one Seurat object (Seurat 4.3.0 ^63,64^) for each brain sample with parameters min.cells = 2 and min.features = 200 (filtered out genes that express in no more than 2 cells, and filtered out cells that have no more than 200 expressed genes). The count matrix from all brains is then integrated together using *RunHarmony* from harmony R package (0.1.1 ^58^) based on STransform processed data and regressed out on factors including individual library and experimental batches.

#### Brain gene expression

To compare gene expression in brain cells, we use the LogNormalize method from the Seurat R package. Raw reads counts were first normalized by the library size and then log-transformed.

#### Single cell clustering and cell type identification

The integrated brain object is then subjected to dimensionality reduction using UMAP methods based on the first 20 principal components from PCA using the Seurat R package. All the cells are then clustered in an unsupervised method using a shared nearest neighbor graph based on k-nearest neighbors (k=20) calculated from the top two coordinates of UMAP. Then the clusters were derived by optimizing the modularity function using the function *FindClusters* with parameters resolution at 0.05. ExN and InN are clustered by first extracting the subset of cells and re-clustering at resolution of 1.

We assigned the cell types to each cluster by the known cell type specific marker gene expression level. Cell type specific marker genes are based on previous publications. For major brain cell types ^37^, subclusters (Azimuth) ^65^, and Vascular cell ^66^. For each cluster, it is assigned to one cell type (A) if two times the average expression of all marker genes of cell type A plus the proportion of cells in the cluster expressing cell type A’s marker genes are larger than any other cell types. We designed a score that takes both the cell type marker gene expression level and proportion of cells expressing the genes into consideration but with a higher emphasis on expression level.

#### Single cell LCS-erosion score and transcriptomic age calculation

We calculated the Local Chromatin Structure (LCS) erosion score for each cell by determining the middle point of the genomic distances of the top 10 most frequently contacting genomic bins. To derive the contact frequency versus genomic distances heatmap for all frontal cortex cells, we first generated 149 genomic bins spanning from 5000 bp to 150M bp with the *n* th bin spanning 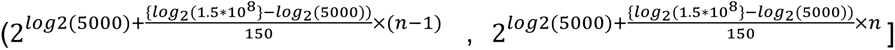 where n=1,2,3,..,150. Subsequently, for DNA reads originating from DD or RD clusters, the genomic distances of all intrachromosomal pairwise interactions were determined, and the frequencies for each genomic distance bin were calculated. These frequencies were then normalized against the bin size. For each single cell the total contact frequency was normalized against the total frequency within each cell before plotting the heatmap. Cells exhibiting an LCS-erosion score exceeding 3e5 were classified as LCS-eroded cells, while others were considered LCS-preserved.

For assessing the transcriptomic-based biological age of cells, we implemented the SCALE methodology described by previous study ^42^. This approach involves using a list of human aging-associated genes obtained from https://sysomics.com/AgingMap/. Initially, we determined the direction (either +1 or −1) of each marker gene based on its correlation with the chronological age of the sample. This direction depended on whether the gene expression was positively (+1) or negatively (−1) correlated with age. Subsequently, for each marker gene, we assigned a weight computed as the proportion of cells expressing the gene, multiplied by its directional value. Finally, we calculated the transcriptomic age for each cell using the dot product of the gene expression z-scores and the corresponding gene weights.

#### LCS-erosion score associated transcriptome functions

To identify genes whose expression is significantly associated with chromatin LCS-erosion score, we applied ANOVA test between each gene’s expression and LCS-erosion score across all cells. Genes with p value < 0.01, F value > 1 and the absolute value of spearman correlation between LCS-erosion score and gene expression larger than 0.1 are considered significant. To identify cell-type specific LCS-erosion score associated genes, we applied ANOVA test within each cell type individually. We used R package *gprofiler2* ^67^ for pathway enrichment analysis. Enriched wikipathway, KEGG and REACtom pathways are shown.

#### eQTL analysis

All brain cell-type specific eQTL data were downloaded from ^44^. We composed a combined dataset by selecting eQTLs with nominal p-value < 10^-4 and removing eQTLs in endothelial cells and pericytes. MUSIC DD contacts between eQTL and their target genes are read pairs where one end is mapped (1-base overlap) over the eQTL and the other end over the gene promoter (2.5 kb flanking region from the TSS). The global Chi-square test was performed on a 6 x 6 table to test the overall association of cell-type specific DD contacts with cell-type specific eQTL-gene pairs. The 6 x 6 table was then reduced to six 2 x 2 tables to test that association in each cell type (Chi-square test). The 95% confidence interval of the odds ratio (OR) was calculated as exp(log(OR) +/-1.96*SELOR), where SELOR is the standard error of the log(OR).

#### XIST-chromatin interactions analysis

To derive the one dimensional XIST-genome attachment level plot (RAL), we extracted all clusters that contained at least one XIST RNA. For each cluster with M XIST RNA reads and N DNA reads, we derived M*N paired XIST-DNA interactions, each of these interactions were then normalized according to their cluster size 1/(M+N). We then binned the whole genome with a 1Mb sized bin. Then the XIST RAL for each bin is the total weights of all XIST-DNA interactions with DNA ends overlapped with that bin.

To derive the 2D RNA-DNA contact map for chrX, we extracted all RNA and DNA reads that can map to chrX. Next, for any matched molecular complex barcode with M chrX RNA reads and N chrX DNA reads, we again first derive all combinations of RNA-DNA interactions with adjusted weight indicating the reverse of cluster size (1/M+N). We binned chrX at 1Mb resolution. The M[i, j] then represents the sum of weighted interactions whose RNA ends mapped to the i th bin and DNA ends mapped to the j th column.

#### XAL stratification and corresponding genomic distance versus contact frequency

XIST-chrX association levels (XAL) is numerically represented as the number of XIST attached chrX DNA bins, each individual cell will have one XAL value. To assess the differences of chromatin organization between XIST+ and XIST-clusters, we stratified all brain cells into four groups based on the XAL value. Group1 with XAL equals 0 will be all cells with no detectable XIST-chrX association. Group2 to Group4, which with increasing XAL value, are taking all cells that have XIST-chrX association and dividing them equally into 3 groups. To derive the genomic distances and contact frequency relationship we follow the same methods introduced in “Calculate genomic distance versus contact frequency curve from MUSIC data ’’ section.

## Acknowledgments

We thank Dr. Thomas Beach and Banner Sun Health Research Institute for providing the human frontal cortex samples and UCSD IGM Genomics Center for support on sequencing.

## Author Contributions

N.C.T. and S.Z. conceived of the MUSIC experimental strategy. Z.L. developed the MUSIC workflow and performed the experiments. X.W. and R.C. performed data analysis. W.Z. X.W. and J.R. contributed to generating data. X.W., Z.L., R.C., W.Z., and S.Z. wrote the manuscript. S.Z. managed and supervised the project.

## Human sample acquisition

The acquisition of postmortem brain is conducted at Banner Sun Health Research Institute with IRB approval (Study Number: 1132516, Investigator: Thomas Beach, M.D., Ph.D.). Informed consent was obtained from all tissue donors.

## EXTENDED DATA

**Extended Data Fig. 1.**
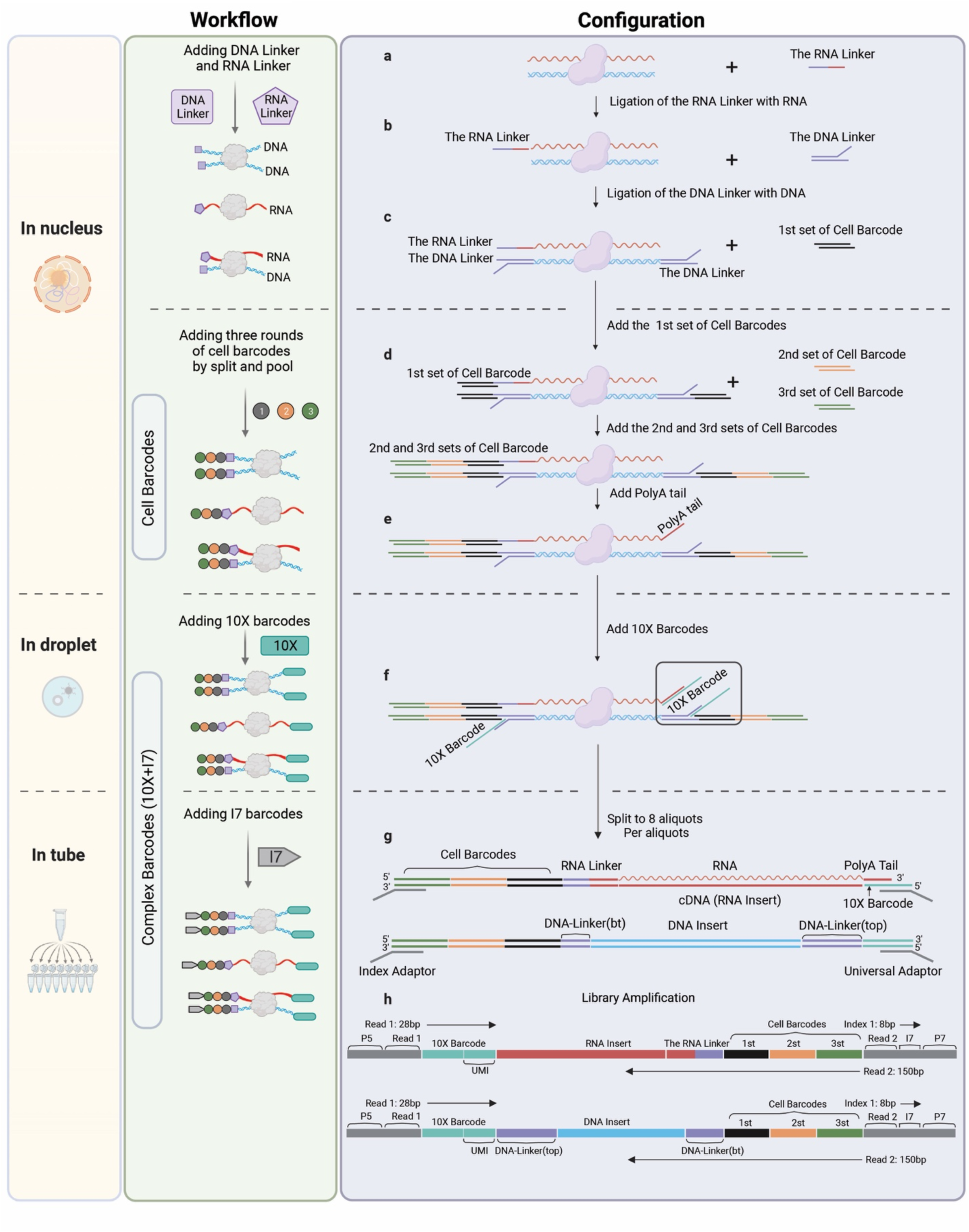
Overview of the MUSIC method. (a) An example chromatin complex with one associated DNA fragment (blue) and one RNA molecule (red) is shown to illustrate the procedure. Other chromatin complexes with any number of associated DNA fragments or RNA molecules are expected to react accordingly. The RNA Linker with half single strand RNA (ssRNA, red) and half single strand DNA (ssDNA, purple). (b) The ligation product with a RNA linker conjugates with the Y-shaped DNA Linker that contains a double strand region (dsDNA) and single strand regions (top-ssDNA and bottom-ssDNA). (c) The ligation product with both DNA and RNA linkers. (d-e) Addition of the three sets of Cell Barcodes (color coded in black, yellow, green) and A-tail. (f) Reaction in a 10X GEM system. (g) Aliquoting the output of the 10X GEM system for library preparation. (h) The sequence configuration of the constructed sequencing library. For Illumina paired-end sequencing, Read 1 covers the 28bp 10X Barcode that consists of a 16bp 10X GEM Barcode and 12bp 10X UMI. Index 1 covers the 8bp I7 Barcode, and Read 2 covers the Cell Barcodes as well as the RNA Linker and the RNA insert or the DNA Linker and the DNA insert.

**Extended Data Fig. 2.**
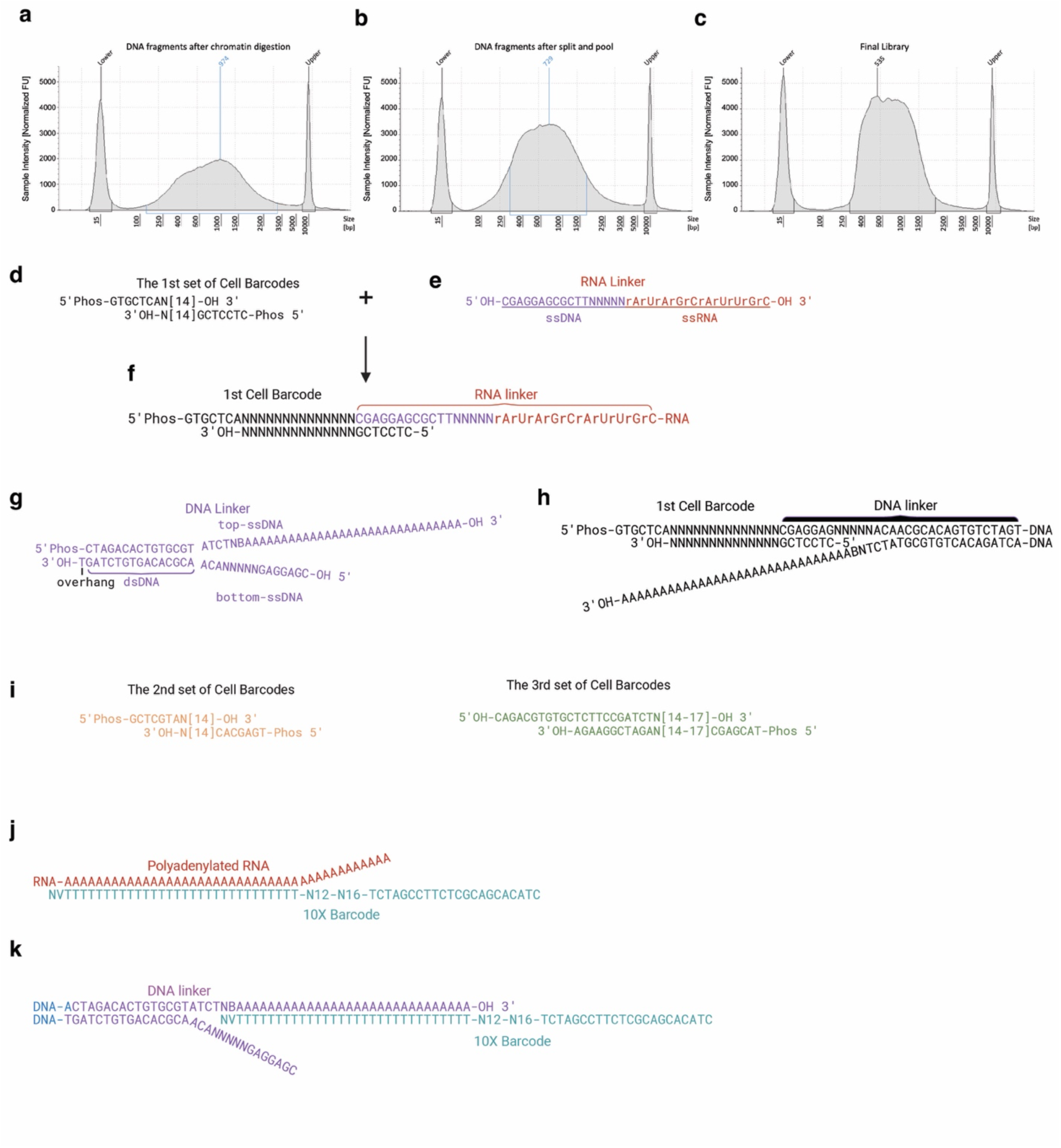
Critical reagents and reactions. (a-c) DNA size distribution of DNA fragments (a) after chromatin digestion, corresponding to Figure 1a, (b) after three rounds of cell barcode ligation, corresponding to Extended Data Fig. 1e, and (c) in the final library, corresponding to Extended Data Fig. 1h. (d) the 1st set of Cell Barcodes. (e) The RNA Linker with half single strand RNA (ssRNA, red) and half single strand DNA (ssDNA, purple) (f) Base-pairing assisted ligation of the ssDNA (purple) in the RNA Linker with the 1st set of Cell Barcodes (black). (g) The Y-shaped DNA Linker with a double strand region (dsDNA) and single strand regions (top-ssDNA and bottom ssDNA) with a 3’dT overhang. (h) The DNA Linker (purple) is ligated with 3’ dA-tailed double-stranded genomic DNA (blue) and with the 1st set of Cell Barcodes (black). (i) the 2nd and the 3rd sets of Cell Barcodes. The 3rd set of Cell Barcodes contains a fraction of Illumina Read2 adapter sequence. (j) Hybridization of the RNA’s polyA tail (red) with the poly(dT) region of the 10X Barcodes (turquoise). (k) Hybridization of the top-ssDNA of the DNA linker (purple) with the poly(dT) region of the 10X Barcodes (turquoise).

**Extended Data Fig. 3.**
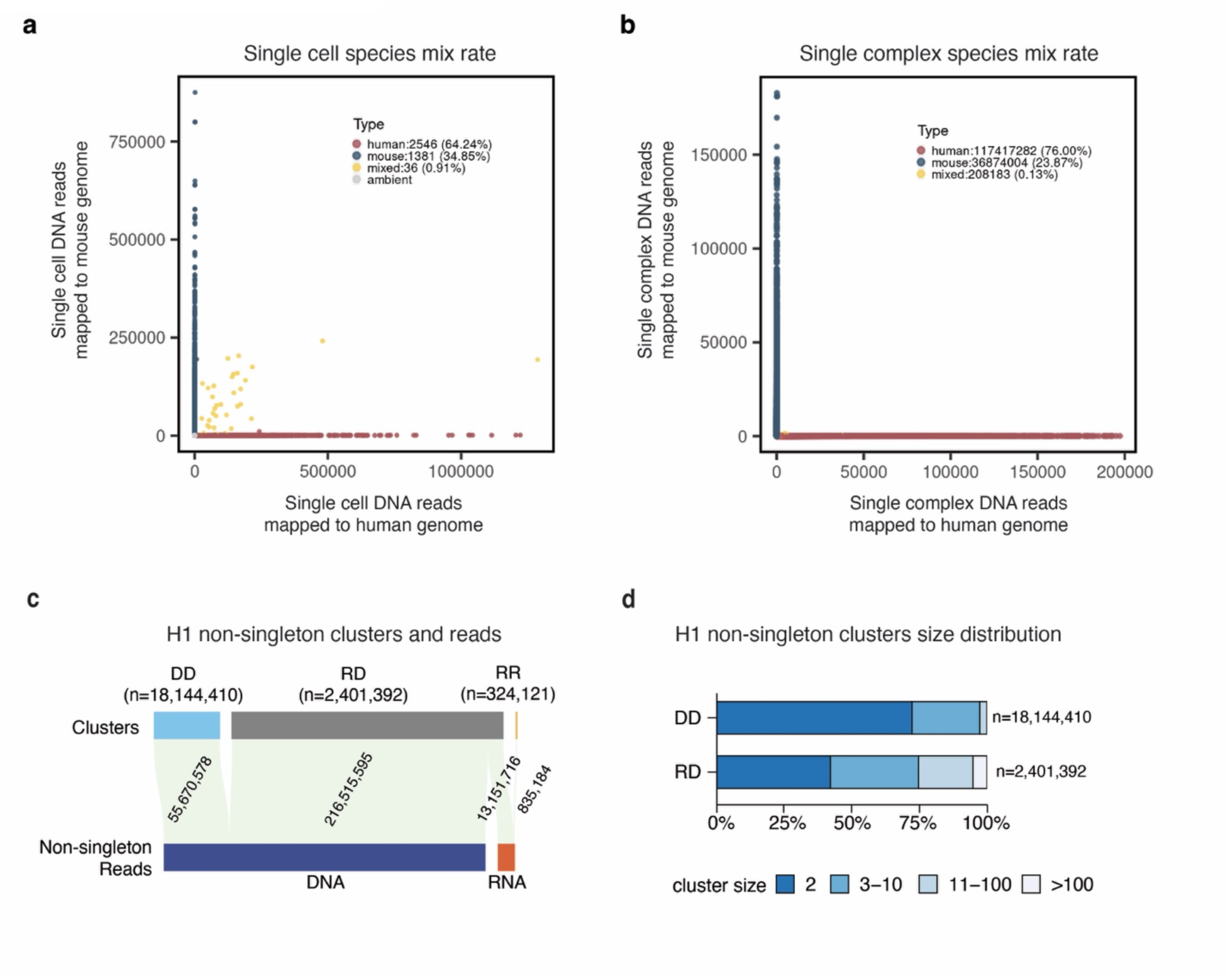
Mixed-species analysis. (a) The cells (dots) with more than 1000 reads (read count filter = 1000) are colored coded to red (human), blue (mouse) if 95% or more of the reads are mapped to a single species (purity filter = 95%), or yellow (mixed) if otherwise. The cells with less than 1000 reads (gray dots, ambient) are not used in the calculation of mixed-species rate. (b) The complexes (dots) are colored coded to red (human), blue (mouse) when 99% or more of the reads are mapped to a single species, or yellow (mixed) if otherwise. (c) The distribution of DNA reads and RNA reads (in gray) in DD, RD, and RR clusters. (d) Cluster size distributions of DD and RD clusters.

**Extended Data Fig. 4.**
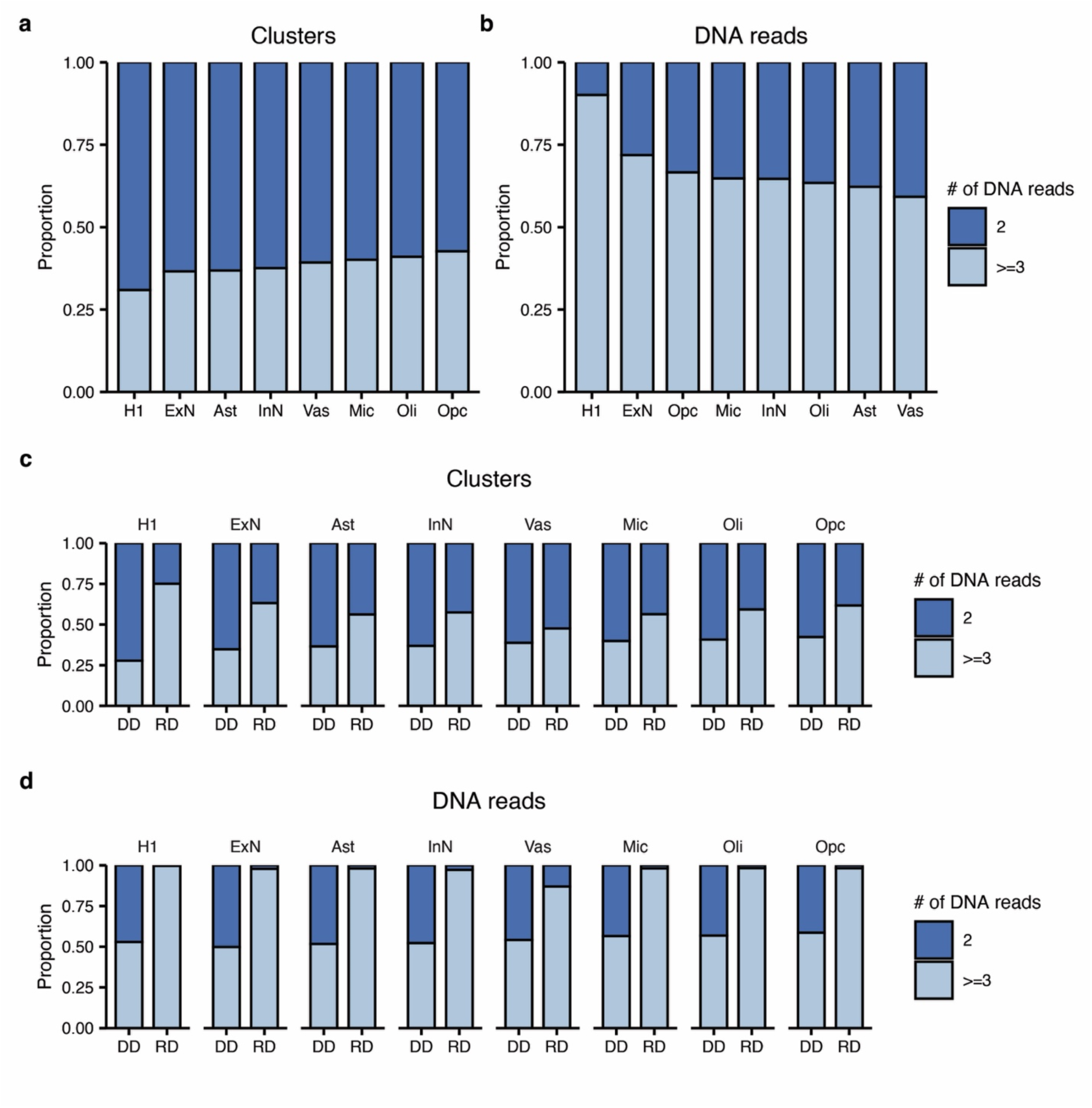
Proportions of pairwise or multiplex interactions. The proportions of clusters (a) and DNA read counts (b) that consist of multiplex chromatin interactions (DNA reads >=3, light blue) in each cell type. Both RD and DD clusters are included in this analysis. (c) Proportions of clusters with pairwise (2 DNA reads, dark blue) or multiplex chromatin interactions (3 or more DNA reads, light blue) in DNA-only (DD) and RNA-DNA (RD) clusters in each cell type. (d) Proportions of DNA read counts with clusters corresponding to pairwise (dark blue) or multiplex (light blue) chromatin interactions.

**Extended Data Fig. 5.**
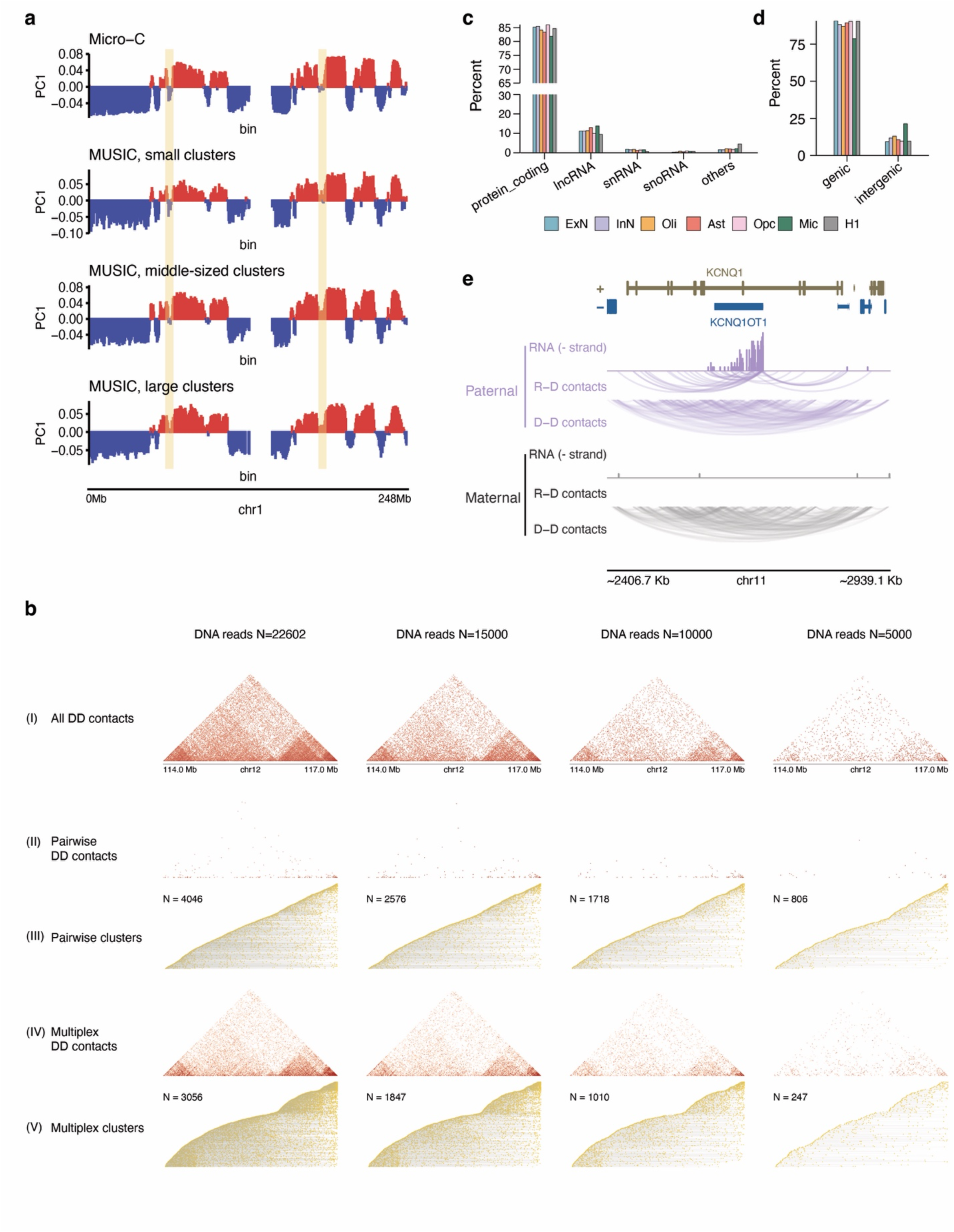
Chromatin contacts and RNA in H1 cells. (a) Compartment scores (PC1) across the entire Chromosome 1 at 500 Kb resolution. PC1s are calculated from Micro-C, MUSIC’s small clusters (2-10 DNA reads), middle-sized clusters (10-50 DNA reads), and large clusters (50-100 DNA reads). Positive and negative PC1s correspond to A and B compartments. Two small B compartments in MUSICs’ small clusters became indistinguishable in the larger clusters (vertical yellow stripes), corresponding to slightly fewer A/B compartment switches in the small clusters. (b) Downsampling the original 22,602 UMNDBC DNA reads mapped to this region (Chr12:114-117Mb) to 15,000, 10,000, and 50,000 reads (columns). Rows from the top to the bottom are 2D contact map of all DD contacts that include multiplex and pairwise contacts (I), 2D contact map of pairwise DD contacts (II), a cumulative view of every DD cluster in pairwise complexes (III), 2D contact map of multiplex DD contacts (IV), and a cumulative view of every DD cluster in multiplex complexes (V). (c) The proportions of each type of RNA species (y axis) in each cell type (color coded). (d) The proportions of genic and intergenic RNA reads in each cell type. (e) Genome track view of the RNA reads (RNA), RNA-DNA (R-D) contacts, and DNA-DNA (D-D) contacts in the paternal (top) and maternal (bottom) allele in H1 cells. All the RNA reads shown are those mapped to the KCNQ1OT1 gene in the strand consistent with the direction of transcription of the KCNQ1OT1 gene (−strand).

**Extended Data Fig. 6.**
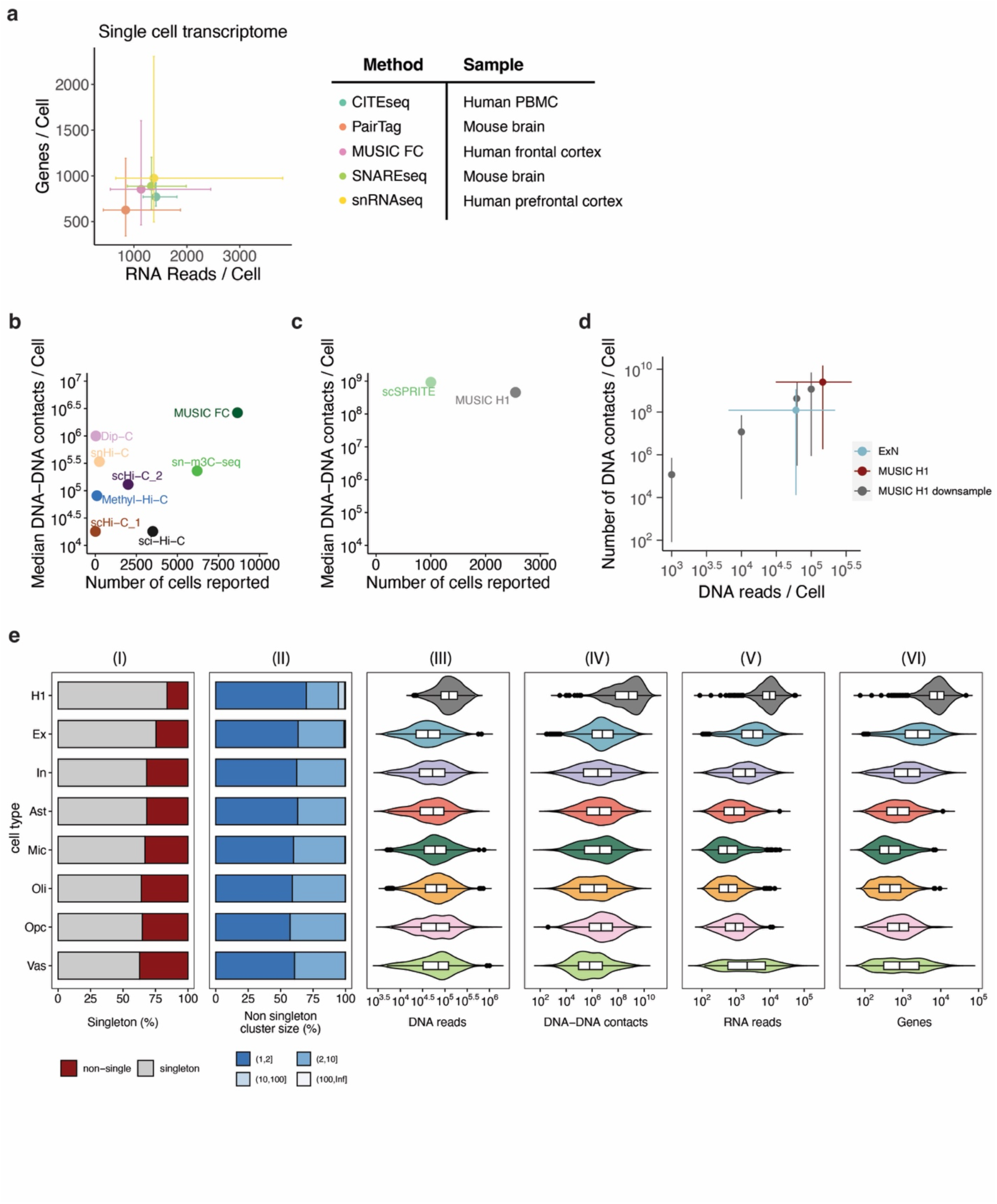
Comparison of technologies. (a) The number of RNA reads per cell (x axis) vs. the number of detected genes per cell (y axis) of CITE-seq, SNARE-seq, PairTag, snRNA-seq, SNARE-seq and MUSIC frontal cortex data (MUSIC FC). The central dot represents the average number and the bars indicate the first and the third quartile of the distribution of all the cells (x axis) and per cell total detected genes (y axis). (b-c) The number of cells (x axis) vs. the median number of DNA-DNA contacts per cell (y axis) for each technique. For MUSIC and scSPRITE, the number of pairwise DNA-DNA contacts are decomposed from multiplex clusters. (d) Downsampling of the sequencing reads of the H1 library. Excitatory neuron (ExN) is used as an example cortical cell type to compare with the downsampled H1 data. The number of chromatin contacts per cell (y axis) is plotted against the number of DNA reads per cell (x axis). The mean (dot) and 95% range of the distribution (whiskers) of the original H1 (red), the downsampled H1 data (gray), and ExN (blue). (e) Summary of MUSIC data by cell type. (I) The proportions of singleton (gray) and non-singleton clusters (red). (II) Cluster sizes of the non-singleton clusters. (III-VI) The distributions of the numbers of DNA reads, DNA-DNA contacts, RNA reads, and detected genes in the single cells. Sample sizes for H1, Ex, In, Ast, Mic, Oli, Opc, Vas are 2267, 1457, 771, 571, 661, 4539, 328 and 760 respectively. For the middle boxplot, the left, center and right edge represent the 25th percentile, median and 75th percentile, respectively. Whiskers extend to 1.5 times the Interquartile Range from the box edges. Data points beyond the whiskers are outliers.

**Extended Data Fig. 7.**
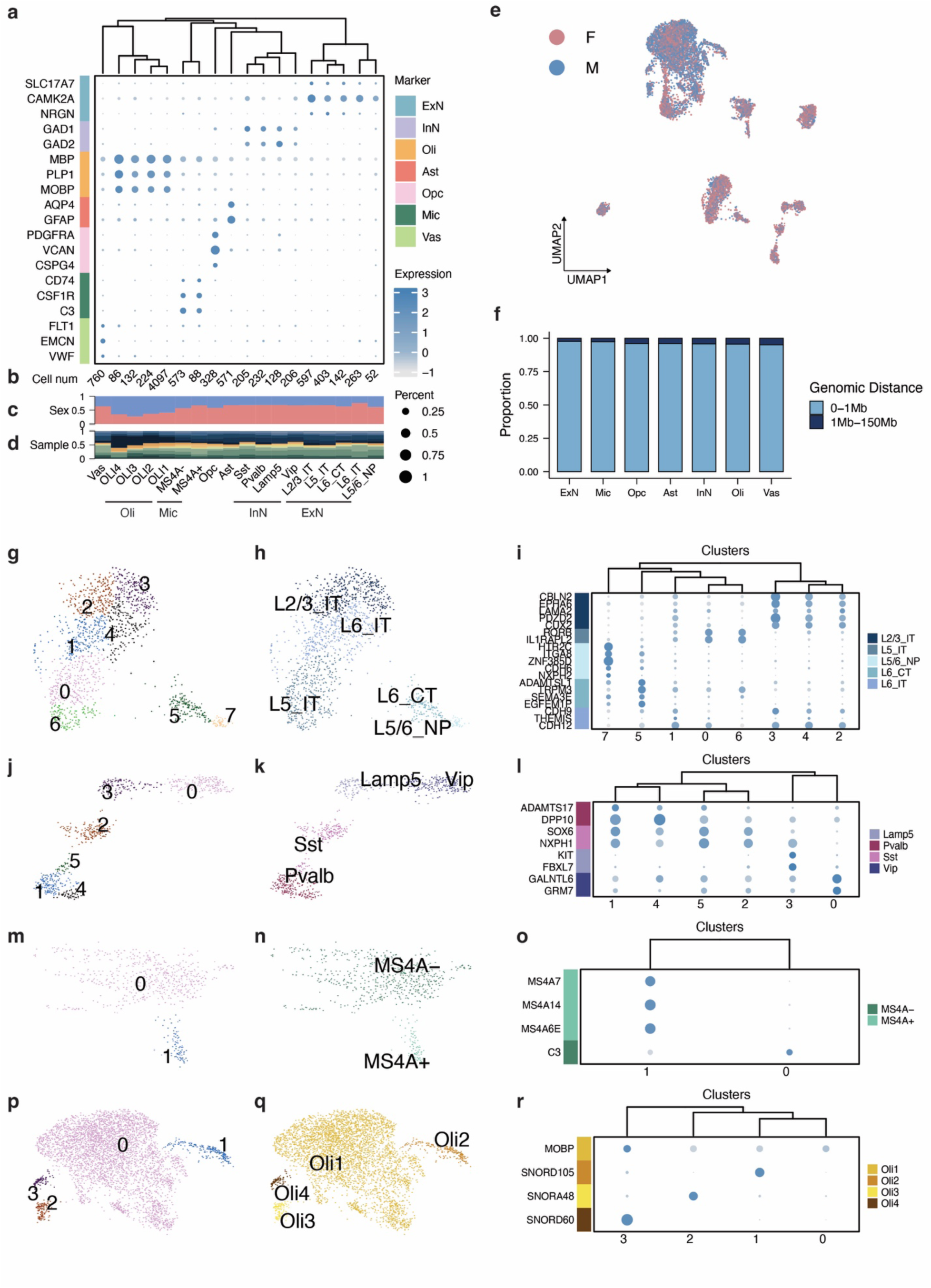
MUSIC analysis of human frontal cortex. Marker gene expression in brains (a), number of cells (b), percent of female (pink) and male cells (blue) (c), and the relative proportion of each sample (d) in every single-cell cluster, and the assigned cell type and subtype for each cluster. Assigned subtypes and marker gene expression in excitatory neurons (g-i), inhibitory neurons (j-l), microglias (m-o), and oligodendrocytes (p-r). (e) Female (pink) and male (blue) cells in the UMAP embedding, showing a sex stratification. (f) Histogram of the proportions of single cells (y axis) with their most frequent chromatin interactions in each genomic bin (0-1Mb, 1Mb-150Mb) (x axis) in each cell type.

**Extended Data Fig. 8.**
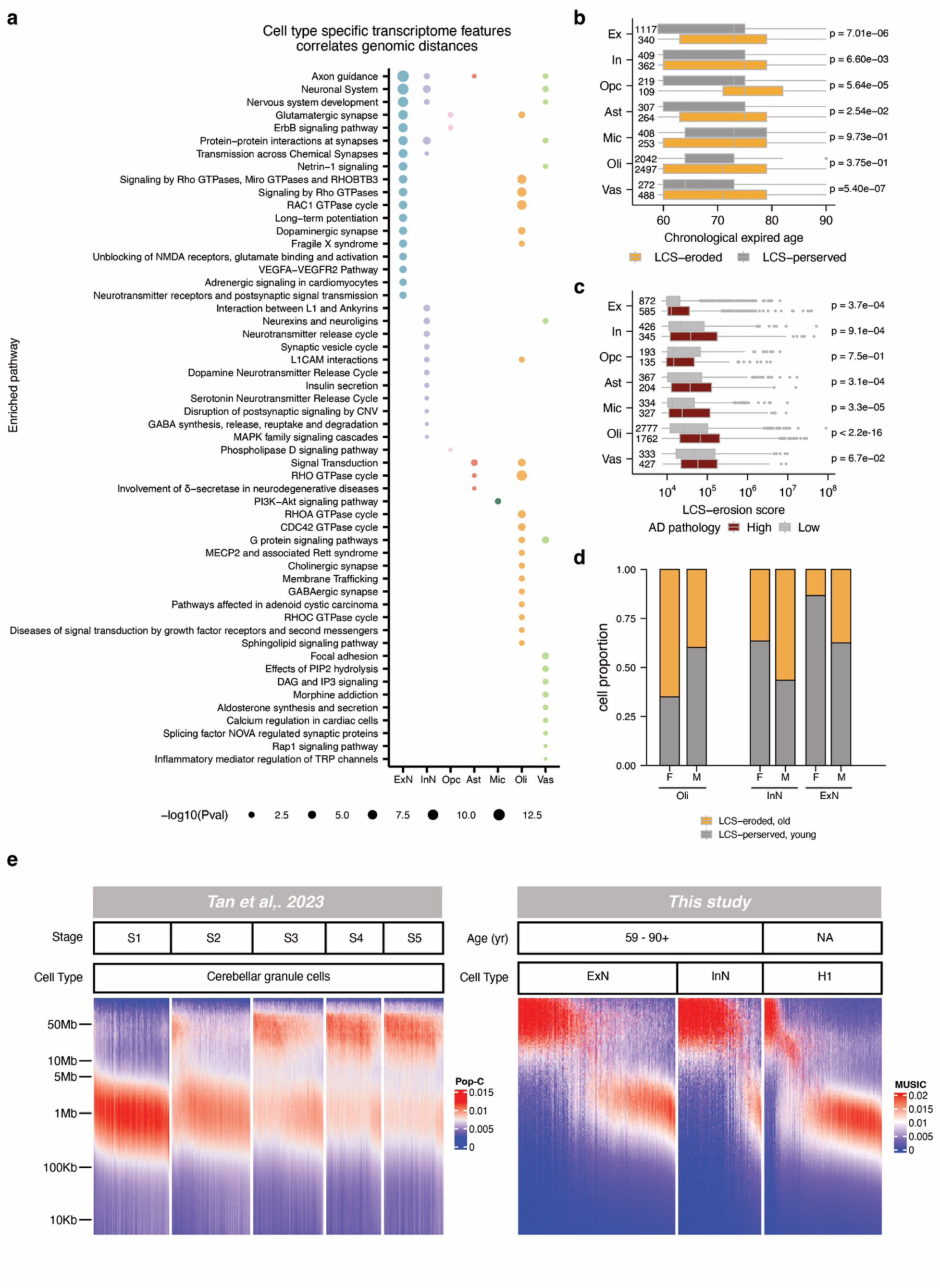
Correlation between Pc(s) and transcriptomic age among single cells. (a) The enriched pathways (row) in those genes that exhibit a correlated expression level with the LCS-erosion score across all the single cells in each cell type (column). The size of the dot represents the enrichment score (−log10(P value)) with larger dots indicating higher significance. (b) Distribution of the chronological ages (age at death, y axis) in the LCS-preserved (gray) and LCS-eroded cells (orange) in each cell type (column). Wilcoxon test, One-sided. Left numbers: sample size. Boxplot: the right edge, center line and left edge represent the 75th percentile, median and 25th percentile, respectively. Whiskers extend to 1.5 times the Interquartile Range from the box edges. Data points beyond the whiskers are outliers. (c) LCS-erosion scores (y axis) in the cells within the samples with high (Braak stage >=4) and low (Braak stage < 3) AD pathology. Wilcoxon test, One-sided. Left numbers: sample size. Boxplot definition is the same as (e). (d) Proportion of LCS-eroded cells and LCS-preserved cells by gender (female (F), male (M)) within oligodendrocytes (Oli) and Neurons (InN, ExN). (e) Number of contacts (color) plotted against genomic distance (y-axis) in single nuclei (columns) using data from Tan et al. (left) and our study (right). Rows represent genomic bins with exponentially increasing sizes. The two datasets were plotted using the same computational program and genomic bin sizes. The ’Stage’ labels correspond to cell groups identified by Tan et al., where Stages S1 to S5 are enriched with cells of chronological ages 0.2, 1, 10, 30, and 80, respectively. ExN: Frontal cortex excitatory neurons. InN: Frontal cortex inhibitory neurons.

**Extended Data Fig. 9.**
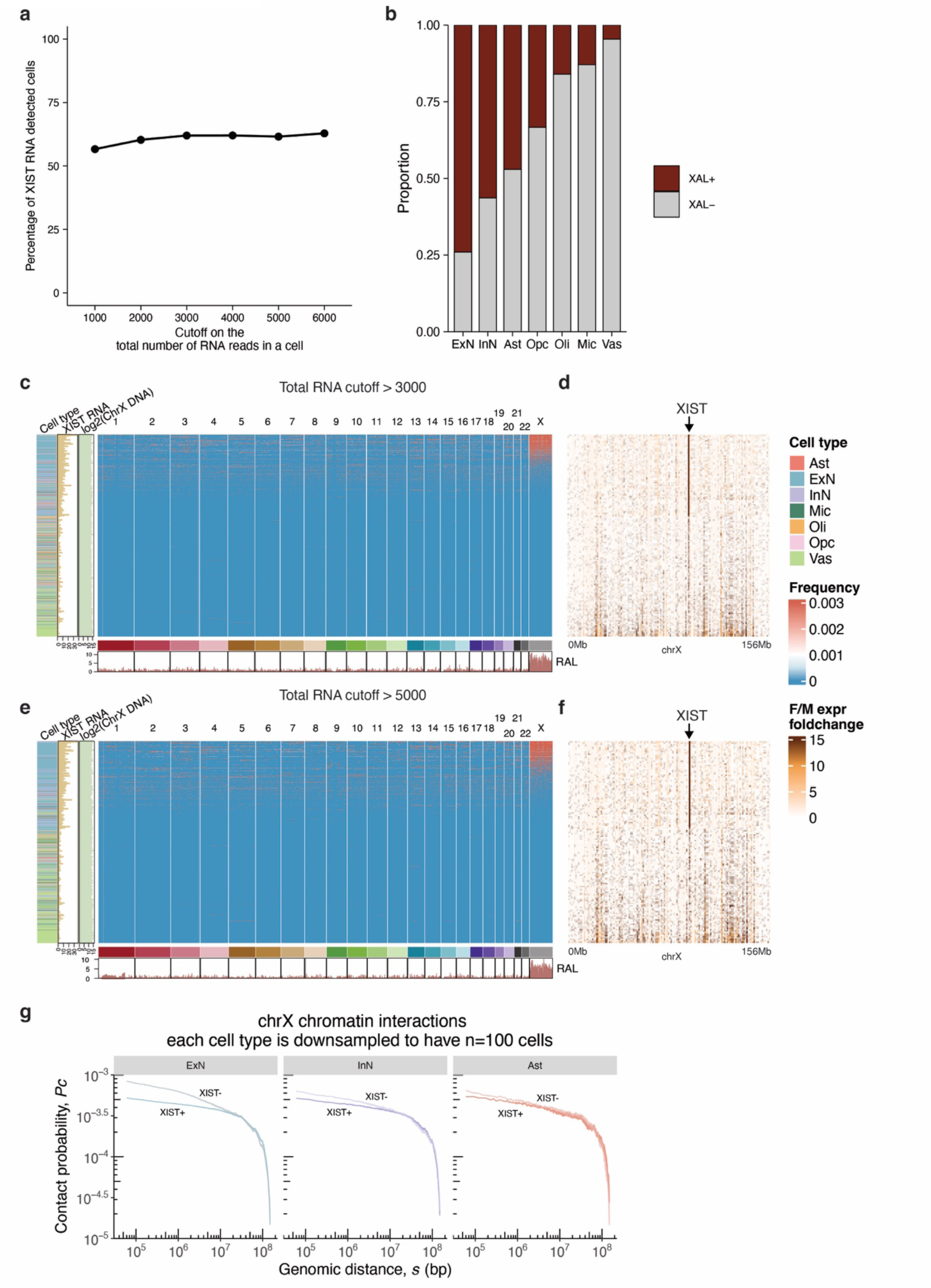
XIST-chromatin association in single female cells. (a) The change of the percentage of XIST lncRNA detected cells (y axis) in the female cortical cells that satisfy the threshold on the total number of RNA reads in a cell (x axis). (b) The variation of the proportions of the observed XAL+ cells across cell types (columns) in the female cortex under total RNA larger than 5000 cutoff. (c-f) The female cells are filtered so that every cell has at least 3000 RNA reads (c-d) or at least 5000 RNA reads (e-f). (c, e) Genome-wide distribution of XIST-chromatin association in individual female cortical cells. Each row represents a single cell. The RNA attachment level (RAL) of the XIST lncRNA (intensity of red color) at any genomic region on any chromosome is plotted with the corresponding genomic coordinates (x axis). Resolution: 1 Mb. Track at the bottom: cumulative RAL of XIST. Tracks on the left indicate the cell type (color), XIST RNA read count (XIST RNA), X chromosomal DNA read counts (ChrX DNA) in log2 scale. (d, f) sex-fold-change of a previously identified gene with incomplete XCI (column) between this female cell (row) and the average expression of the male cells of the matched cell type. Only the genes expressed in at least 20 cells are plotted. (g) Downsampling analysis of cell numbers. X chromosome Pc(s) curves in excitatory neurons (ExN), inhibitory neurons (InN), and astrocytes (Ast) are plotted from the same number of female cells (n=100). The difference between XIST+ (darker color) and XIST-Pc(s) (lighter color) curves is most pronounced in ExN.

