## Supplementary material for "Single-cell multiplex chromatin and RNA interactions in aging human brain": Full supplementary file

**Table of Contents**


Supplemental Note 1 2
Supplemental Note 2 2
Supplemental Note 3 4
Supplemental Note 4 5
Supplemental Note 5 6
Supplemental Note 6 6
Supplemental Note 7 6
Supplemental Note 8 8
Supplemental Note 9 8
Supplemental Note 10 9
Supplemental Note 11 10
Supplemental Note 12 10
Supplemental Note 13 11
Supplemental Note 14 11
Supplementary Table 1 14
Supplementary Table 2 15
Supplementary Table 3 16
Supplementary Table 4 17
Supplementary Table 5 18
Supplementary Table 6 19
Supplementary Fig. 1. DNA size distribution and nuclei isolation. 20
Supplementary Fig. 2. Clustering analysis of the integrated dataset. 21
Supplementary Fig. 3. Cell type specific chromatin contacts and eQTL gene interaction. 22
Reference 23

### Supplemental Note 1

Single-cell multi-omic technologies have revolutionized our ability to collectively analyze epigenomic or chromatin conformation features and gene expression [1–5]. Various methodologies have been introduced to enable the joint profiling of specific aspects of the genome, transcriptome, or proteome. For instance, SNARE-seq [2] and SHARE-seq [4] allow for simultaneously profiling chromatin-accessible regions (CARs) and gene expression (Supplementary Table 1). Additionally, scNMT-seq [6] facilitates the combined analysis of CARs and DNA methylation, while scNOMeRe-seq [7] enables the joint examination of CARs, DNA methylation, and gene expression. Techniques such as Pair-tag provide insights into histone modification profiles and transcription levels [8], and sn-m3C-seq captures chromatin interactions and DNA methylation patterns [9]. CITE-seq, on the other hand, allows for the simultaneous measurement of surface protein and transcriptome [10]. Recently, HiRES technology has been introduced to the concurrent application of Hi-C and RNA-seq within individual cells [5] (Supplementary Table 1). Furthermore, imaging tools like DNA seqFISH+ [11], ORCA [12], and MINA [13] have facilitated the joint analysis of chromatin traces and gene expression (Supplementary Table 1).

### Supplemental Note 2

The MUSIC workflow contains two major steps.

#### Step 1: Adding Cell Barcodes

The first step ligates *the RNA Linker* to the RNA molecules and *the DNA Linker* to the fragmented DNA and adds the Cell Barcodes (Extended Data Fig. 1a-e). This step accomplishes the first and the second design goals. The RNA Linker is a chimeric oligonucleotide consisting of single-stranded DNA (ssDNA) at its 5' end and single-stranded RNA (ssRNA) at its 3' end (Extended Data Fig. 2e). The ssRNA and ssDNA are designed to facilitate efficient ligations with RNA and the Cell Barcodes, respectively. The ligation of the RNA Linker with the RNA produces a chimeric product “RNA-Linker+RNA” with ssDNA on this 5’ end (Extended Data Fig. 1b).

The DNA Linker is partially double-stranded DNA (dsDNA) and partially single-stranded DNA (ssDNA). It is generated by hybridizing two DNA strands (Extended Data Fig. 2g, Supplementary Table 2). The DNA Linker is designed to (1) efficiently ligate with fragmented chromosomal DNA through the dsDNA region via sticky-end ligation, (2) effectively ligate with the Cell Barcodes via the ssDNA region on the bottom strand, and (3) facilitate hybridization with the 10X Barcodes through the ssDNA region on the top strand. Ligation of the DNA Linker with fragmented DNA generates the DNA-Linker+DNA+DNA-Linker structure.

Subsequently, both RNA-Linker+RNA and DNA-Linker+DNA+DNA-Linker undergo ligation with the Cell Barcodes, which label the RNA and DNA originating from the same nuclei. An innovation in the MUSIC workflow is the ability to employ the same procedure for adding Cell Barcodes to both RNA and DNA. The ssDNA region on the 5' end of the RNA Linker

(RNA-Linker_ssDNA) and the ssDNA region on the bottom strand of the DNA Linker (DNA-Linker_bt-ssDNA) share the same optimized DNA sequence, enabling efficient ligation with the Cell Barcodes.

The Cell Barcodes consist of three sets: the 1st, 2nd, and 3rd sets of Cell Barcodes (Extended Data Fig. 2d, i). Each set comprises a top-strand overhang, a dsDNA region, and a bottom-strand overhang. The top-strand overhang and bottom-strand overhang are identical for all barcodes within a set, while the dsDNA region is unique for each barcode. The addition of the three sets of barcodes is accomplished through three rounds of split-pooling of the nuclei [14] (Extended Data Fig. 1d-e).

The three sets of Cell Barcodes are designed to maximize the ligation efficiency for sequentially ligating the 1st set of Cell Barcodes with the RNA Linker and the DNA Linker, the 2nd set with the 1st set, and the 3rd set with the 2nd set. The optimal ligation efficiency is achieved by the complementarity of the 1st set's overhang with RNA-Linker_ssDNA and DNA-Linker_bt-ssDNA and the complementarity of the overhang sequences of the sequentially ordered barcode sets (Extended Data Fig. 1c, Extended Data Fig. 2f, h). Out-of-order ligations, such as the ligation of the 2nd set of Cell Barcodes with the RNA Link are minimized due to lack of sequence complementarity in the overhangs by design [15,16].

**Step 2: Adding Complex Barcodes**

The second step of the MUSIC workflow involves pooling the nuclei and releasing the molecular complexes from the nuclei. At this stage, the molecular complexes can be traced back to individual nuclei using their Cell Barcodes. The next objective is to assign the same Complex Barcode to any RNA or DNA within the same molecular complex. The combination of Cell Barcodes and Complex Barcodes specifies each molecular complex in a particular cell. It should be noted that the Complex Barcodes alone, without the Cell Barcodes, are not expected to be sufficient for identifying every molecular complex.

The Complex Barcode comprises two sets of barcodes: the 10X Barcodes and the I7 Barcodes that are embedded in the Index Adaptors, a.k.a. I7 Barcodes. We chose to use the combination of 10X Barcodes and I7 Barcodes rather than the 10X Barcodes alone as the Complex Barcodes because the combination offers an X-fold increase in the number of barcodes than the 10X Barcodes alone, where X is the number of distinct I7 Barcode sequences.

The 10X Genomics' Gel Bead-in-Emulsion (GEM) system is employed to add the 10X Barcodes. This system encapsulates molecular complexes into individual GEMs, also called droplets, where the complexes in each droplet are added with a unique 10X Barcode. Importantly, the MUSIC technology by design does <u>not</u> require most droplets to encapsulate no more than one molecular complex. As we will explain later, a single molecular complex is identified by the combination of Cell Barcodes and Complex Barcodes. This droplet-based addition of 10X Barcodes can be considered another round of split-pooling, where the units are molecular complexes, unlike the previous rounds where the units were nuclei.

The addition of 10X Barcodes is required for both RNA and DNA. The 10X system is optimized to add 10X Barcodes to RNA. The RNA, following polyadenylation, can be hybridized by the polyT sequence within the 10X Barcodes (Extended Data Fig. 1f), initiating subsequent automated cDNA synthesis within each GEM. In the case of DNA, MUSIC leverages the design of the DNA-Linker to add 10X Barcodes. The DNA-Linker contains a 30 nt single-stranded polyA sequence (Extended Data Fig. 2g), which can hybridize with the 30 nt PolyT sequence in the 10X Barcodes (Extended Data Fig. 2k). This allows subsequent DNA synthesis using the fragmented DNA as a template.

The 10X GEM system comprises approximately 3.5 million unique 10X Barcodes. This number exceeds the expected number of molecules captured per cell by a single-cell assay. Consequently, there is no significant concern regarding the sufficiency of 10X Barcodes to differentiate molecular complexes that can be captured and assayed from a single cell. However, including I7 Barcodes in combination with the 10X Barcodes ensures the sufficient diversity of Complex Barcodes.

The I7 Barcode, a part of the Index Adaptor, is added during the PCR process that constructs the final sequencing library. The output of the 10X GEM system comprises two components: cDNA originating from RNA and double-stranded DNA (dsDNA) originating from chromosomal DNA. The cDNA takes the form of Cell Barcode+RNA-Linker+RNA Insert+10X Barcode, while the dsDNA takes the form of Cell Barcode+DNA-Linker(bt)+DNA Insert+DNA-Linker(top)+10X Barcode. DNA-Linker(top) and DNA-Linker(bt) denote the dsDNA originating from the top and the bottom strands of the DNA Linker. Both the cDNA and dsDNA contain the 3rd set of the Cell Barcode at one end, which includes a fraction of the Illumina Read2 sequence, and the 10X Barcode at the other end, including a fraction of the Illumina Read1 sequence. These components can be constructed into the Illumina sequencing library using PCR. The PCR primers, namely the Index Adaptor and the Universal Adaptor, common to both cDNA and dsDNA, are used to amplify them into the final sequencing library.

The addition of the different I7 Barcodes is achieved through another round of split-pooling. The output of the 10X GEM system is split into eight tubes. Each tube is added with an Index Adaptor that contains a specific I7 Barcode, for PCR-based library construction. The constructed sequencing libraries are combined to form the final sequencing library. In this round of split-pooling, the units are molecular complexes. To summarize, MUSIC employs two rounds of split-pooling to add the Complex Barcodes. The first round involves splitting by 10X GEMs, while the second round involves splitting by tubes. In the current implementation (MUSIC v1.0), the estimated number of possible barcode combinations is up to 25 trillion, approximated by multiplying the 884,736 distinct Cell Barcodes with 3.5 million available 10X Barcodes, and further multiplying this total by the 8 different I7 Barcodes.

**Supplemental Note 3**

We used the mix-species dataset to evaluate MUSIC's resolution at single-cell and single-complex levels. The mixed-species dataset resolved 372,878,969 DNA reads and

22,714,916 RNA reads for humans (hg38), 128,687,584 DNA reads and 8,951,899 RNA reads for mice (mm10). All the subsequent analyses are based on these UMNDBC reads.

First, MUSIC identifies the DNA reads and RNA reads that share the same Cell Barcodes (CB) as coming from the same cell. Based on an established approach and threshold [14], MUSIC resolved 2,546 human cells, 1,381 mouse cells, and 36 cells with mixed species content. This resulted in a species-mixing rate of 0.91% at the cell level (Extended Data Fig. 3a). Comparatively, this species-mixing rate is within the same order of magnitude as the species-mixing rate of 3% reported for scSPRITE under the same thresholds [14]. Next, MUSIC identifies the reads sharing the Cell Barcodes and the Complex Barcodes (CB+10X+I7) as a cluster. In the mixed-species dataset, 154,499,469 clusters were resolved, of which 154,291,286 clusters contained reads from a single species. This corresponds to a complex-level species-mixing rate of 0.13% (Extended Data Fig. 3b). Taken together, the low species-mixing rates at the cell level and the complex level support the ability of MUSIC to generate data at single-cell and single-complex resolutions.

We analyzed the human H1 cells regarding the numbers of reads, clusters, and pairwise contacts. First, the data solved 2,546 human H1 cells. Each H1 cell contained an average of 144,049 UMNDBC DNA reads (blue dots) and 11,384 UMNDBC RNA reads (orange dots) (Figure 1b). Second, clusters were defined as collections of reads sharing the same Cell Barcode and Complex Barcode (CB+10X+I7). In the perfect scenario where cluster-level species-mixing rate = 0, a cluster reflects a molecular complex. A cluster is called a DNA-only, an RNA-only, or an RNA-DNA cluster if it contains only DNA reads, only RNA reads, or both DNA and RNA reads. On average, each H1 cell had 7,036 DNA-only clusters (DD clusters), 232 RNA-only clusters (RR clusters), and 1,170 RNA-DNA clusters (RD clusters) (Figure 1c). Finally, co-complexed multi-way associations were projected to co-complexed read pairs based on a previously described procedure [14], resulting in an average of 2,639,302,084 co-complexed DNA-DNA pairs, 7,089,720 co-complexed RNA-RNA pairs, and 250,525,581 co-complexed RNA-DNA pairs per cell (Figure 1d).

**Supplemental Note 4**

We carried out a downsampling analysis to test whether the different read numbers in pairwise and multiplex interactions can explain the better representation of TAD structure in the multiplex interactions. We downsampled the total number of 22,602 UMNDBC DNA reads in this region (Chr12:114-117Mb) (complete data) to 15k, 10k, and 5k DNA reads (downsampled datasets). At the extreme end of the downsampled datasets (5k reads, pairwise and multiplex combined), the TAD structure becomes indiscernible, and accordingly there is no discernable TAD structure in either the pairwise or the multiplex clusters (Extended Data Fig. 5b). In the other downsampled datasets (15k, 10k), the TAD structure remained discernable in the total DD contacts (pairwise and multiplex combined). Similar to what's observed in the complete data (without downsampling), pairwise clusters alone poorly reflected the TAD structure, while multiplex clusters recapitulated the TAD structure (Extended Data Fig. 5b). Notably, the numbers of DNA reads in multiplex clusters in these downsampled datasets (7,717 multiplex DNA reads when downsampled to 15k; 3,782 multiplex DNA reads when downsampled to 10k) are smaller than the number of DNA reads in pairwise clusters in the complete data (8,092 pairwise DNA reads), and in contrast, the TADs structures are discernable the multiplex clusters in the downsampled datasets but not in the pairwise clusters in the complete data, suggesting that the stronger representation of TADs in the multiplex clusters is not due to a larger number of DNA reads in the multiplex clusters.

**Supplemental Note 5**

Data from bulk assays revealed that the probability of chromatin interactions (Pc) decreases as the genomic distance (s) between the two DNA fragments increases [17,18]. Furthermore, Pc(s) almost linearly correlates with s at the log-log scale [17,18] (Figure 2e). In MUSIC data, the Pc(s) curve derived from size-2 DD clusters exhibited a linear relationship at the log-log scale (size=2 curve, Figure 2e), consistent with proximity-ligation-based assays. The DD clusters of size 3 or greater also exhibit decreasing Pc(s) curves, which are no longer linear at the log-log scale. The larger the cluster size, the greater Pc(s) deviates from a straight line (Figure 2e). The multiplex interactions displayed higher contact frequencies at submegabase to several megabase genomic distances as compared to the pairwise contacts, suggesting an enrichment of long-range chromatin interactions in the multiplex complexes.

**Supplemental Note 6**

Previous bulk assays studies have identified two key features of genome-wide RNA-chromatin associations. First, genomic regions with high levels of locally transcribed premature mRNA (pre-mRNA) (pre-mRNA-rich regions) tend to correspond to the A compartments defined by chromatin conformation assays such as Hi-C and Micro-C [19,20]. Second, these pre-mRNA-rich regions also serve as hotspots for long-range RNA-chromatin interactions involving non-coding RNAs MALAT1, 7SK, and snRNAs (collectively called nsaRNAs) [19]. In the MUSIC-identified RNA-DNA contacts (ensemble RD), the pre-mRNA associated genomic regions correlated with Micro-C derived A compartment, and more specifically with the “Speckle” nuclear compartmentalization jointly inferred from TSA-seq, DamID, and Hi-C data [21] (Figure 2l). To categorize an RNA read as pre-mRNA, we required it to span an intron-exon junction and contain at least 15 intronic nucleotides. Furthermore, the genomic regions targeted by long-range nsaRNA-chromatin associations (blue curve) overlapped with regions associated with high levels of locally transcribed pre-mRNA (p-value < 2.2e-16, red curve, Figure 2l, m). Thus, MUSIC recapitulated both prominent features of genome-wide RNA-chromatin associations. In addition, the MUSIC data reproduced the allele-specific association of the KCNQ1OT1 noncoding RNA with the genomic regions adjacent to the KCNQ1OT1 gene [22] (Extended Data Fig. 5e), affirming MUSIC's ability to recapitulate the known chromatin association pattern of a specific RNA.

**Supplemental Note 7**

We compared the number of RNA reads per cell between MUSIC and other technologies. We compared the RNA reads in MUSIC FC to single-nucleus RNA-seq data from human prefrontal cortex (snRNA-seq PFC) [23] and multi-omic datasets including CITE-seq data generated from human PBMC [10], SNARE-seq data generated from mouse brain [2], and PairTag data generated from mouse brain [1] (Supplementary Table 5). snRNA-seq PFC resolved a median of 1,973 RNA reads per nucleus, corresponding to 1,348 genes. CITE-seq, SNARE-seq, and PairTag resolved a median of 1416, 1332 and 845 RNA reads per nucleus, corresponding to 770, 1203, and 626 genes (Extended Data Fig. 6a). MUSIC FC exhibited a median of 1,136 RNA reads per nucleus, corresponding to 853 genes, which was within the range of the other methods (Extended Data Fig. 6b). Thus, MUSIC demonstrated a comparable read depth to snRNA-seq and other multi-omic methods in mapping single-nucleus transcriptomes.

We also compared the numbers of resolved nuclei and DNA-DNA contacts. We compared MUSIC to published single-cell multi-omic datasets of scHiC_1[24], sci-Hi-C[25], scHi-C_2[26], snHi-C[27], Dip-C[28], scSPRITE[14] and multi-omics methods Methyl-Hi-C[29], sn-m3C-seq[9]. Most of these datasets contained up to 2,000 nuclei, while the sn-m3C-seq dataset had 6,500 nuclei (Extended Data Fig. 6b, c). We reused the numbers of median DNA-DNA contacts per cell of each technology from a recent review[30]. MUSIC FC had 9,087 nuclei and a median of 2,651,577 DNA-DNA contacts, surpassing most other datasets (Extended Data Fig. 6b, c). Moreover, scSPRITE data generated from mouse embryonic stem cell (mESC) [14] resolved marginally fewer nuclei and more DNA-DNA contacts, both within 2-fold of difference as compared to MUSIC H1 data (Extended Data Fig. 6c).

We compared the usable fractions of reads between MUSIC and scSPRITE. The scSRIPTE mESC dataset includes a total number of 1,269,693,929 raw sequencing reads, of which 632,242,064 (49.8%) contains all the six barcodes [14]. scSPRITE has 50.2% loss due to incomplete cell barcodes, which is greater than MUSIC's total loss of Cell Barcodes and Complex Barcodes (35.94%). After read trimming, mapping, and removing PCR duplicates, the final yield of scSPRITE is 161,989,473 reads, which account for 12.75% of the raw sequencing reads. In comparison, MUSIC’s final yield (UMNDBC reads) account for 17.39% of the raw sequencing reads. Thus, the usable fraction of the total sequencing reads in MUSIC is comparable, if not higher, than in scSPRITE. Taken together, these data suggest comparable throughputs between MUSIC and the other technologies. Of note, previous work projected scSRITE’s multiplex interactions to pairwise contacts and reported the projected pairwise contacts as the number of DNA-DNA contacts of scSPRITE [14]. We followed this approach to project MUSIC’s multiplex data to pairwise contacts. The greater numbers reported here for scSPRITE and MUSIC are attributable to their ability to capture multi-way contacts, which does not mean that they are more sensitive in revealing pairwise chromatin interactions than the other methods.

A downsampling analysis suggests that differences in sequencing depth partly account for the higher contact numbers observed in MUSIC H1 compared to MUSIC FC (Extended Data Fig. 6d). Additionally, we acknowledge the potential influences of other biological or technical factors, such as the documented decrease in chromatin contacts during ES cell differentiation [31,32], on these discrepancies.

**Supplemental Note 8**

We performed clustering analysis on the 9,087 cortical nuclei based on their RNA expression levels (Figure 3a). The clusters were then assigned to specific cell types using known marker genes from previous studies (Supplementary Table 6) [23,33]. A cluster of 1,457 nuclei was categorized as excitatory neurons (ExN) based on the expression of marker genes SLC17A7, CAMK2A, and NRGN [23] (Extended Data Fig. 7a). Within this cluster, two subclusters were identified: intratelencephalic (IT) neurons (on the left) and corticothalamic (CT) and near-projecting (NP) neurons (on the right) (Extended Data Fig. 7g-i). The IT subcluster further consisted of three subgroups: Layer 2/3 intratelencephalic (L2/3 IT) (597 nuclei), Layer 5 IT (403 nuclei), and Layer 6 IT (263 nuclei) neurons (Extended Data Fig. 7g-i). The CT and NP subcluster contained Layer 6 corticothalamic (L6 CT) (142 nuclei) and Layer 5/6 near-projecting (L5/6 NP) (52 nuclei) neurons, respectively [34] (Extended Data Fig. 7g-i).

A cluster of 771 nuclei was categorized as inhibitory neurons (InN) based on the expression of marker genes GAD1 and GAD2 [23] (Extended Data Fig. 7a). This cluster formed four subclusters (Extended Data Fig. 7j-l) corresponding to Lamp5, Vip, Pvalb, and Sst neurons. The two sub-clusters on the top corresponded to Lamp5 (128 nuclei) and Vip (206 nuclei) neurons that develop from caudal ganglionic eminence [35]. The two sub-clusters on the bottom corresponded to Pvalb (232 nuclei) and Sst (205 nuclei) neurons that develop from medial ganglionic eminence (Extended Data Fig. 7j-l).

A cluster of 661 nuclei was categorized as microglia based on the expression of marker genes CD74, CSF1R, and C3 [23] (Extended Data Fig. 7a). This microglial cluster consisted of a major subcluster (573 nuclei) and a minor subcluster (88 nuclei) marked by different levels of Membrane-spanning 4A (MS4A) genes [36] (Extended Data Fig. 7m-o). In mice, the activation of MS4A genes marks the transition of microglia from a chemokine state to an interferon state [36]. Thus, the major and the minor sub-clusters may reflect the corresponding cellular states of microglia in humans.

A cluster of 4,539 nuclei was categorized as oligodendrocytes based on the expression of marker genes MBP, PLP1, and MOBP [23], which included a major subcluster (4,097 nuclei) and three minor subclusters (224, 132, and 86 nuclei) (Extended Data Fig. 7p-r). Another cluster of 760 nuclei was categorized as vascular cells based on the expression of marker genes EMCN, VWF, and FLT1 [33] (Figure 3a). Lastly, two distinct clusters with 571 and 328 nuclei were categorized as astrocytes (Ast) based on marker genes AQP4 and GFAP, and oligodendrocyte precursors (Opc) based on marker genes VCAN, PDGFRA, and CSPG4 [23]. Additionally, a joint analysis of MUSIC FC data with a single-nucleus RNA-seq (snRNA-seq) dataset of human frontal cortex revealed highly consistent clustering structures and clustering-based cell type assignments between the two datasets (Supplemental Note 9).

**Supplemental Note 9**

We evaluated the consistency of single-cell clustering and cell type assignments between MUSIC FC and a snRNA-seq dataset of human frontal cortex [23] (hereafter called Mathys' dataset). We integrated the two datasets using the Harmony software [37] and carried out Uniform Manifold Approximation and Projection for Dimension Reduction (UMAP) analysis on the integrated dataset (Supplementary Fig. 2a-c). The single cells in the integrated dataset exhibited clear clustering structure, and the co-clustered single cells originated from MUSIC FC and Mathys' dataset tended to share the same marker gene expression profiles (Supplementary Fig. 2d-g), suggesting a high degree of consistency between MUSIC FC and Mathys' dataset.

MUSIC FC's oligodendrocyte cluster included three previously uncharacterized sub-clusters. These sub-clusters (Oli2, Oli, Oli4) are separated from the main oligodendrocyte cluster (Oli1) due to their cluster-specific expression of snoRNAs (Extended Data Fig. 7r). snoRNAs are non-polyA RNAs that cannot be captured by the the snRNA-seq technique used by Mathys et al., whereas they are included in MUSIC FC because MUSIC captures both polyA and non-polyA RNA types (Extended Data Fig. 5c). Accordingly, we expect these oligodendrocyte sub-clusters to be grouped in the main oligodendrocyte cluster in the integrated dataset, which indeed was the case (Supplementary Fig. 3d-g). Furthermore, 76.62%, 73.21%, 68.18% and 81.39% cells from MUSIC FC's Oli1 (the main cluster), Oli2, Oli3, and Oli4 were included in the oligodendrocyte cluster of the integrated dataset, revealing a high degree of consistency between Oli2, Oli3, Oli4 and the oligodendrocyte cluster of the integrated dataset.

**Supplemental Note 10**

We analyzed the contributions of transcriptomic age, chronological age, sex, AD pathology (AD or non-AD), and cell type to explain the variation of LCS erosion score. To ensure the robustness of the results, we ran two linear models, namely a full model (Full Model) and a model selection identified model (Selected Model). The two models led to essentially the same results.

First, we specified the Full Model as:
$log(Y) = \beta_0 + \beta_1 X_1 + \beta_2 X_2 + \beta_3 X_3 + \beta_4 X_4 + \beta_5 X_5 + \beta_{6-31}(X_1 * X_2 * X_3 * X_4 * X_5) + \varepsilon$ , where Y denote the LCS erosion score, $X_1$ denotes Transcriptomic age, $X_2$ denotes Chronological Age, $X_3$ denotes AD pathology, $X_4$ denotes Sex, $X_5$ denotes Cell type, and $X_1 * X_2 * X_3 * X_4 * X_5$ denotes the collection of all the pairwise and higher order interaction terms. The covariates collectively explain 27.25% of the variance in $log(Y)$. Six terms exhibit the ability to explain 1.0% or more variance, which are:

1. Transcriptomic age, explaining 14.38% of the variance (p value < 2.2e-16),
2. Sex * Cell type, explaining 2.09% of the variance (p value < 2.2e-16),
3. Cell type, explaining 1.83% of the variance (p value < 2.2e-16),
4. Sex, explaining 1.67% of the variance (p value < 2.2e-16),
5. AD pathology, explaining 1.57% of the variance (p value < 2.2e-16),

6. Transcriptomic age * Cell type, explaining 1.10% of variance (p value < 2.2e-16),

None of the other covariates explains 1% of the total variance.

Next, we identified the Selected Model using stepwise model selection with the optimal Akaike information criterion (AIC), as:

$log(Y) = \beta_0 + \beta_1 X_1 + \beta_3 X_3 + \beta_4 X_4 + \beta_5 X_5 + \beta_7 X_1 X_3 + \beta_9 X_1 X_5 + \beta_{15} X_4 X_5 + \varepsilon$ , i.e.

$$log(LCS\ erosion\ score) \sim \beta_1 Transcriptomic\ age + \beta_3 Pathology + \beta_4 Sex + \beta_5 Cell\ type$$

$$+ \beta_7 Transcriptomic\ age * Pathology + \beta_9 Transcriptomic\ age * Cell\ type + \beta_{15} Sex * Cell\ type$$

The Selected Model explains 23.34% of the variance in $log(Y)$, where Transcriptomic age, Sex*Cell Type, Cell type, AD pathology, Sex, Transcripomic age*Cell type, and Transcriptomic age*Pathology explains 14.38%, 2.65%, 2.23%, 1.70%, 1.15%, 0.72%, 0.52%, respectively. This Selected Model includes all the top six terms with the largest explanatory abilities in the Full Model, suggesting the two models prioritized the same explanatory variables. In summary, LCS erosion score correlates with transcriptomic age, cell type, AD pathology, sex and the interactions of sex and cell type, transcriptomic age and cell type, and transcriptomic age and pathology.

**Supplemental Note 11**

MUSIC did not detect XIST lncRNA in every female cell. There are two probable causes for this variation. Technically, MUSIC may not be sensitive enough to detect XIST lncRNA in every nucleus. Biologically, the expression level of XIST may be heterogeneous among the female cells. Supporting the coexistence of both causes, XIST lncRNA is exclusively detected in female cells, and as the threshold on the number of total RNA reads in a cell is increased, the proportion of XIST-detected cells among the remaining female cells remains relatively stable with a modestly increasing trend (Extended Data Fig. 9a). This analysis suggests that the RNA read count in a cell is influenced by both the expression level of the molecule and the sensitivity of the technique itself. To mitigate the impact of false negatives in our analyses, we refined the dataset by filtering based on RNA read count, and furthermore, we refrained from drawing conclusions about individual cells.

**Supplemental Note 12**

We compared XIST-chrX association with chrX gene expression. Without a phased genome, MUSIC data cannot infer allelic expression and thus cannot directly pinpoint any gene expressed from the Xi chromosome (incomplete XCI gene). However, a survey conducted by the Genotype-Tissue Expression (GTEx) consortium, involving 5,500 transcriptomes from 449 individuals across 29 tissues, reported that incomplete XCI genes generally exhibit higher expression in females compared to males in the corresponding tissue, suggesting sex bias can be used as “a proxy for XCI status" [38]. The MUSIC data can infer sex-difference in gene expression at the single-cell level. To illustrate this point, for every female cell and every previously annotated incomplete XCI gene [38], we calculated the fold change between the RNA read count of this gene in this female cell and the average RNA read counts of this gene in all the male cells of the matching cell type (sex-fold-change) (Extended Data Fig. 9d, f). XAL- female cells exhibited greater sex-fold-changes in most of the incomplete XCI genes compared to XAL+ female cells, suggesting the loss of XIST-chrX association in single female cells correlates with larger sex-difference in the gene expression in the human cortex (Extended Data Fig. 9d, f).

**Supplemental Note 13**

Aligned with our findings, seqFISH+ analysis in the female mouse cortex revealed that Xa has smaller 3D spatial distances than Xi when the genomic sequence is below ~10Mb, and vice versa when the genomic distance is above ~10Mb (Fig. S24g in [39]). Moreover, the female mouse cortex demonstrated a clearer difference between Xa and Xi in ExN compared to InN and Ast (Fig. S24 in [39]). Specifically, most single ExN cells exhibited a smaller intra-chromosomal spatial distance in Xi than Xa at genomic distances greater than ~10Mb. In contrast, a notable fraction of InN and Ast cells showed non-distinguishable differences in intra-chromosomal spatial distance between Xi and Xa at any genomic distance (Fig. S24g in [39]).

**Supplemental Note 14**

We developed a data processing pipeline, called MUSIC-docker for passing the MUSIC sequencing data. We packaged the data processing snakemake pipeline [59] with the Docker platform to enable execution of this data processing pipeline in any operating system without the need for re-compilation of the source codes. MUSIC-docker takes in i7 index splitted pair-end fastq files and process them into two sequence files (.fastq), corresponding to the RNA inserts and the DNA inserts, based on the RNA Linker or the DNA-Linker(bt) sequence identified in each read pair. MUSIC-docker adds the Cell Barcodes, the 10X Barcode, and the I7 Barcode into the sequence name of each RNA or DNA sequence. MUSIC maps the RNA sequences and the DNA sequences to the genome to obtain two mapped files (.BAM), where every sequence is associated with its genomic coordinate and with its Cell Barcodes and Complex Barcodes (see Methods). A full documentation of the MUSIC-docker pipeline is available at: http://sysbiocomp.ucsd.edu/public/wenxingzhao/MUSIC_docker/intro.html.

Convert the raw sequencing output to FASTQ files

The sequencing output file (.bcl) is provided together with the eight I7 Barcode sequences to the *bcl2fastq* program to yield eight FASTQ files. Each FASTQ file contains read pairs with a 28bp Read1 and a 150bp Read2. The 28bp Read1 sequence contains the 10X Barcode, which is composed of a 16bp 10X GEM Barcode and a 12bp 10X UMI sequence. The 150bp Read2 sequence contains the 3rd, 2nd, 1st Cell Barcodes, the RNA Linker sequence or the DNA-Linker(bt) sequence, the RNA insert or the DNA insert.

Extracting Cell Barcodes

Demultiplexing is the step to extract the fragment identity information (FMI) and the fragment sequence information (insert) from paired-end FASTQ files and reformat them into two new FASTQ files for RNA and DNA reads individually where the read sequence will be the insert, and the read name will be the fragment identity information. This step is applied individually to each I7 index separated library.

FMI includes three cell barcodes, 10X GEM barcodes and 10X UMI. The 3rd Cell Barcode (CB3) contains a 14-17bp variable-length unique sequence region, followed by a 7bp overhang sequence, while the 2nd (CB2) and the 1st (CB1) Cell Barcodes contain a 14bp unique sequence region, followed by 7bp overhangs. CB3 has varied length to introduce complexity of the base composition at the same position which will help improve the sequencer's base calling quality. The linker sequences followed by the Cell Barcodes in Read2 are either the RNA Linker or the DNA-Linker(bt). The RNA Linker contains a 7bp overhang-complementary sequence to CB1 and a 20bp sequence that is unique to the RNA Linker. We note that RNA Linker's 7bp overhang has been counted in CB1's overhang sequence. The DNA-Linker(bt) is the double-stranded DNA produced from the bottom strand of the DNA Linker. The DNA-Linker(bt) is 31bp in length, including a 7bp overhang sequence and a 24bp sequence that is unique to the DNA Linker. We note that the 7bp overhang sequence has been counted in CB1's overhang. Based on whether it is the RNA Linker or the DNA-Linker(bt) sequence, we separately output to two .fastq files.

Following the RNA Linker sequence is the RNA insert, which is output as the sequence of the .fastq file for the RNA. The output read name for this RNA sequence is in the format of [previous read name]|[CB3]-[CB2]-[CB1]-[10X GEM Barcode]-[I7 Barcode]#[10X UMI], where [I7 Barcode] is a choice of "lib1", …, "lib8". The following is an example output sequence.

**@MN00185**:343:000H532W5:1:11102:14470:1060|BC3_68-BC2_89-BC1_46-TGACGCGGTACACGTT-lib1*#GTAA G GGTGAAG*

AATGGCAGAATGTTTTGGTCAAAATTTTTTAGCCAGCACCCCAATTAAAAAAAAAAAAAAAAA
A +

FAFFFF==AFFFFFAFAFFAAF6=AFFFFFFA=F=/FAF=FFF=A=FFFF=FFFFF=/FF=//6

Following the DNA-Linker(bt) sequence is the DNA insert, which is output as the sequence of the .fastq file for the DNA. The output read name for this DNA sequence is in the format of [previous read name]|[CB3]-[CB2]-[CB1]-[10X GEM Barcode]-[I7 Barcode]#[10X UMI], where "I7 Barcode" is a choice of "lib1", …, "lib8". The following is an example output sequence.

**@MN00185**:343:000H532W5:1:11102:15055:1047|BC3_75-BC2_51-BC1_11-CGGAACCAGGCATCGA-lib1*#GGTC T GTACCTG*

TCATAAAGAACTCTACCAACTTAAGAAGAAGAAAGCAAGCAACCCCATACAAAAATGGCG
A +

=F/FA=/F=/F=FF/FF///6/A=AAFFF/A/=//F///FF/FFFFF/FF///FFF/F///

The Cell Barcodes are designed such that the Levenshtein distance between any two of the Cell Barcodes is greater than 2. For Cell Barcode or Linker sequences, we allow at most 2 mismatches (Hamming distance) to the reference sequence. The whitelist of the "10X Single Cell 3' v3.1 Chemistry" - is provided for matching the 10X Barcodes, where a match is defined as a perfect match. The demultiplexing step is implemented in python with customized code.

Reads cleaning

Based on our experimental design and sequencing errors from the Illumina machine, it is possible that we introduced some artificial sequences into the library even if the reads could have perfectly matched the three cell barcodes and 10X barcodes. For DNA reads, we will remove the consecutive As or consecutive Gs from the 3' of the DNA insert if there are more than 20 consecutive As or consecutive Gs. For RNA reads, we will detect the ssDNA region of the RNA Linker sequence "CGAGGAGCGCTT" and remove any sequence following it. We use *cutadaptor* (v2.8) with parameter `-q 15 -m 20` to ensure the remaining sequences have a base quality score larger than 15 and with length at least 20bp.

**Mapping DNA and RNA ends**

After raw reads demultiplexing and reads cleaning, we will map DNA and RNA inserts to the reference genome. We used *bowtie2* (v5.4.0) for DNA inserts genome mapping with the command "bowtie2 -p 10 -t --phred33 -x ". We used *bwa mem* to map RNA inserts to the reference genome in a splice aware manner with parameters "-SP5M". Only uniquely mapped reads were selected and will be used for downstream analysis.

**Reads deduplication**

Following read mapping, the subsequent crucial task involves the removal of PCR duplicates, which are identified by sharing the same Unique Molecular Identifier (UMI). In our study, we utilized the 10X UMI. Deduplication based on mapped coordinates is a widely employed approach, as alignment software typically accounts for sequencing errors present in the reads. To adapt to our specific requirements, we have developed a customized deduplication tool capable of accommodating the customized read name format. The deduplication process involves initially sorting the BAM file according to the mapping coordinates. Subsequently, we perform a single pass scan of the BAM file. Any read that satisfies all the following criteria is flagged as a duplicate: 1) it maps to a location within 8 base pairs of the previous read, 2) it shares the same cell barcode and molecular barcode with the previous read, and 3) its UMI exhibits a Levenshtein distance of less than 2 base pairs from the UMI of the previous read. All identified PCR duplicates are subsequently removed to ensure the integrity of downstream analyses.

**Merging I7 index separated libraries**

In the final step, we consolidate all the deduplicated BAM files obtained from each I7 index-marked library into a unified, comprehensive BAM file. This final output is generated in a sorted BAM format, which captures essential information including the cell barcode, molecular barcode (comprising the 10X and I7 index), and the mapping location of the insert for both DNA and RNA (distinctly stored in separate files).

**Supplementary Table 1.**

**A partial list of single-cell multi-modal technologies.** “CARs”: chromatin accessible regions. “Co-complex pairs”: variations of Hi-C technologies that sample a pair of DNA fragments from a chromatin complex involving either pairwise or multiplex interactions. “Multiplex interactions”: technologies that can measure multiplex interactions and thus differentiate pairwise and multiplex interactions.

| Method | Imaging/Omics | Surface protein | CARs | Histone modification | DNA methylation | 3D genome organization | | | Gene expression | RNA-chromatin association |
|---|---|---|---|---|---|---|---|---|---|---|
| | | | | | | Chromatin traces | Co-complex pairs | Multiplex interactions | | |
| **DNA seqFISH+** | Imaging | | | ✓ | | ✓ | | | ✓ | |
| **ORCA** | Imaging | | | | | ✓ | | | ✓ | |
| **MINA** | Imaging | | | | | ✓ | | | ✓ | |
| **Snare-seq** | Omics | | ✓ | | | | | | ✓ | |
| **SHARE-seq** | Omics | | ✓ | | | | | | ✓ | |
| **scNMT-seq** | Omics | | ✓ | | ✓ | | | | | |
| **scNOMeRe-seq** | Omics | | ✓ | | ✓ | | | | ✓ | |
| **Pair-tag** | Omics | | | ✓ | | | | | ✓ | |
| **sn-m3C-seq** | Omics | | | | ✓ | | ✓ | | | |
| **CITE-seq** | Omics | ✓ | | | | | | | ✓ | |
| **HiRES** | Omics | | | | | | ✓ | | ✓ | |
| **MUSIC** | Omics | | | | | | | ✓ | ✓ | ✓ |

**Supplementary Table 2.**

**Sequences of the RNA Linker, the DNA Linker, Universal adaptor, Index adaptors, and Cell Barcodes.**
https://docs.google.com/spreadsheets/d/1SiIrvWFqIWpHL9TDgs2Rd1RxiE08vFvq9kHXLxWHEOc/edit?usp=sharing

**Supplementary Table 3.**

**Number of reads retained in every data processing step in the mixed-species library.** (b) Imperfect_10x indicates the number of MUSIC sequencing reads that do not have the 16bp 10x barcode sequence that can perfectly align to the whitelist provided by 10X genomics. (c) Incomplete_CB: number of reads that don't have three cell barcode sequences that can map to the cell barcodes whitelist each within 2 edit distances. (h, i) DNA or RNA reads that are uniquely mapped to the human genome. (j, k) PCR duplicates removed uniquely mapped barcode-complete reads. Details of sequencing reads processing steps can be found in the Methods section.

| Steps | Reads Number | Ratio |
|---|---|---|
| (a) Raw_reads | 3,067,956,666 | |
| (b) Imperfect_10x | 101,139,708 | (b)/(a) 3.30% |
| (c) Incomplete_CB | 1,001,280,905 | (c)/(a) 32.64% |
| (d) Adaptor_removed_DNA | 9,591,538 | (d)/(a) 0.31% |
| (e) Adaptor_removed_RNA | 29,416,328 | (e)/(a) 0.96% |
| (f) Barcode complete, adaptor removed DNA reads | 1,291,977,026 | (f)/(a) 42.11% |
| (g) Barcode complete, adaptor removed RNA reads | 195,044,878 | (g)/(a) 6.36% |
| (h) DNA uniquely mapped reads | 732,019,497 | (h)/(a) 23.86% |
| (i) RNA uniquely mapped reads | 45,719,121 | (i)/(a) 1.50% |
| (j) DNA uniquely mapped, non-duplicate, and barcode-complete reads (UMNDBC) | 501,566,553 | (j)/(a) 16.35% |
| (k) RNA uniquely mapped, non-duplicate, and barcode-complete reads (UMNDBC) | 31,666,815 | (k)/(a) 1.03% |

### Supplementary Table 4.

### Brain sample metadata.

| Sample ID | Case ID | Sex | Age at death | ApoE | PlaqueTotal | TangleTotal | Braak score |
|---|---|---|---|---|---|---|---|
| B1 | 12-49 | M | 63 | 3/4 | 14.5 | 15 | VI |
| B2 | 11-38 | M | 64 | 3/3 | 14.5 | 15 | VI |
| B3 | 18-78 | M | 71 | 3/3 | 0 | 4 | III |
| B4 | 13-40 | M | 73 | 3/3 | 0 | 2.25 | II |
| B5 | 19-40 | F | 75 | 3/4 | 15 | 15 | VI |
| B6 | 13-49 | F | 75 | 3/3 | 0 | 2.5 | II |
| B7 | 17-60 | F | 82 | 3/3 | 15 | 14.5 | VI |
| B8 | 15-46 | F | 82 | 2/3 | 1 | 1.5 | I |
| B9 | 11-107 | M | 71 | 4/4 | 14 | 15 | VI |
| B10 | 16-32 | M | 79 | 2/3 | 0 | 1 | II |
| B11 | 10-22 | F | 59 | 3/3 | 1 | 0.5 | I |
| B12 | 09-46 | F | 60 | 3/3 | 15 | 15 | VI |
| B13 | 16-18 | M | ≥90 | 2/4 | 14.5 | 14 | VI |
| B14 | 10-26 | F | ≥90 | 3/3 | 0 | 6.5 | III |

**Supplementary Table 5.**

**Transcriptome profiling statistics from multi-modal single cell sequencing technologies.** All statistics for each technique are computed based on the raw count matrix downloaded from each published dataset (CITE-seq: GSE100866_PBMC, SNARE-seq: GSE126074_AdBrainCortex, PairTag: GSE152020, snRNA-seq: Synapse: syn18485175).

| Method | Median RNA reads / cell | Medium genes / cell | Number of cells in Raw count table | Number of genes in Raw count table | Sample |
|---|---|---|---|---|---|
| **snRNA-seq** | 1,973 | 1,348 | 80,660 | 18,192 | Human prefrontal cortex |
| **CITE-seq** | 1,416 | 770 | 7,985 | 29,929 | Human PBMC cells |
| **SNARE-seq** | 1,332 | 887 | 10,309 | 33,160 | Mouse adult brain cerebral cortex |
| **Pair-Tag** | 845 | 626 | 64,849 | 52,781 | Mouse frontal cortex and hippocampus |
| **MUSIC H1** | 6,559 | 3,684 | 2,546 | 39,007 | Human embryonic H1 cell line |
| **MUSIC FC** | 1,136 | 853 | 9,067 | 22,373 | Frozen human frontal cortex |

### Supplementary Table 6.

### Marker genes for each cell type and subtype.

| Type | Marker genes | Subtype | Marker genes |
|---|---|---|---|
| **ExN** | SLC17A7, CAMK2A, NRGN | L2/3_IT | CBLN2, EPHA6, LAMA2, PDZD2, CUX2 |
| | | L5_IT | RORB, IL1RAPL2 |
| | | L6_IT | CDH9, THEMIS, CDH12 |
| | | L5/6_NP | NPSR1.AS1, HTR2C, ITGA8, ZNF385D, CDH6, NXPH2 |
| | | L6_CT | ADAMTSL1, TRPM3, SEMA3E, EGFEM1P |
| **InN** | GAD1, GAD2 | Vip | GALNTL6, GRM7 |
| | | Lamp5 | KIT, FBXL7 |
| | | Sst | SOX6, NXPH1 |
| | | Pvalb | ADAMTS17, DPP10 |
| **Mic** | CD74, CSF1R, C3 | MS4A- | N/A |
| | | MS4A+ | MS4A7, MS4A14, MS4A6E |
| **Oli** | MBP, PLP1, MOBP | | |
| **Opc** | PDGFRA, VCAN, CSPG4 | | |
| **Ast** | AQP4, GFAP | | |
| **Vas** | FLT1, EMCN, VWF | | |

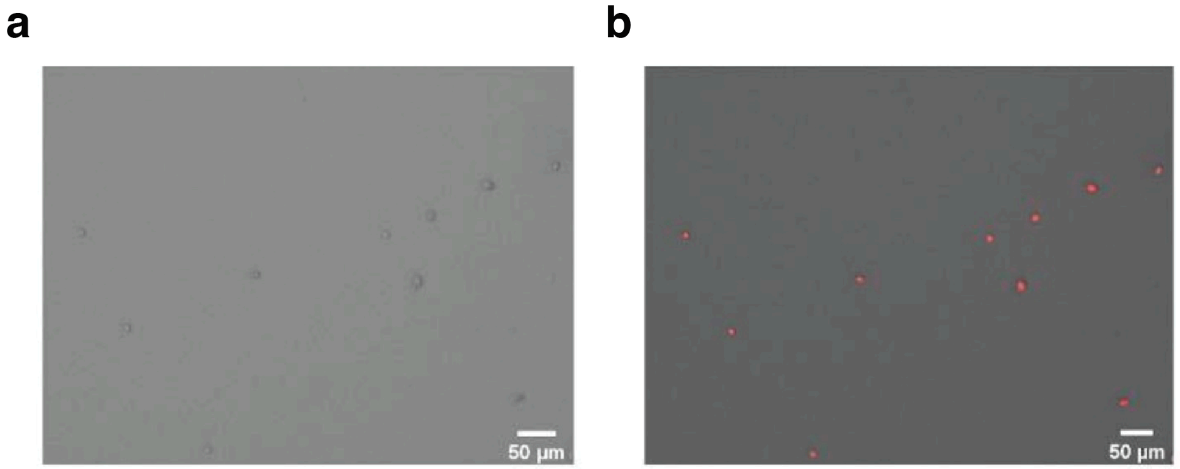


**Supplementary Fig. 1. DNA size distribution and nuclei isolation.**

Isolated nuclei from human cortex under a bright field (a) or fluorescence microscope with EthD-1 staining (red) (b). This experiment was repeated independently three times with similar results.

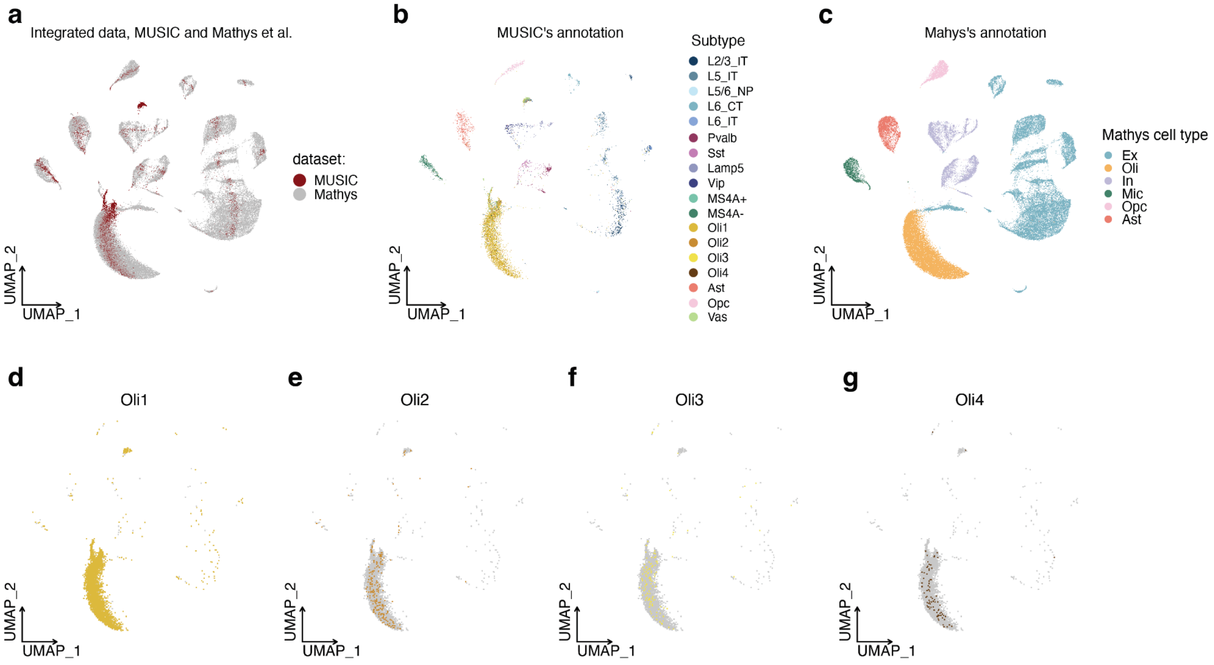


**Supplementary Fig. 2. Clustering analysis of the integrated dataset.**

The integrated dataset is composed of MUSIC FC data and a frontal cortex snRNA-seq dataset from Mathys et al. (a-c) UMAP of integrated MUSIC and snRNA-seq data, with MUSIC cells marked by red and snRNA-seq cells by gray (a), colored by MUSIC's cell type annotation (b), or Mathys et al.'s cell type annotation (c). (d-g) Distribution of MUSIC's oligodendrocyte cluster in the integrated dataset. All the cells in MUSIC's oligodendrocyte cluster are plotted (light gray) and the cells in each sub-cluster (Oli1 - Oli4) are highlighted with color. Most cells in every oligodendrocyte sub-cluster in the MUSIC dataset are within the oligodendrocyte cluster of the integrated dataset, suggesting all the MUSIC identified oligodendrocyte sub-clusters are aligned with an oligodendrocyte cluster from Mathys et al.'s snRNA-seq data.

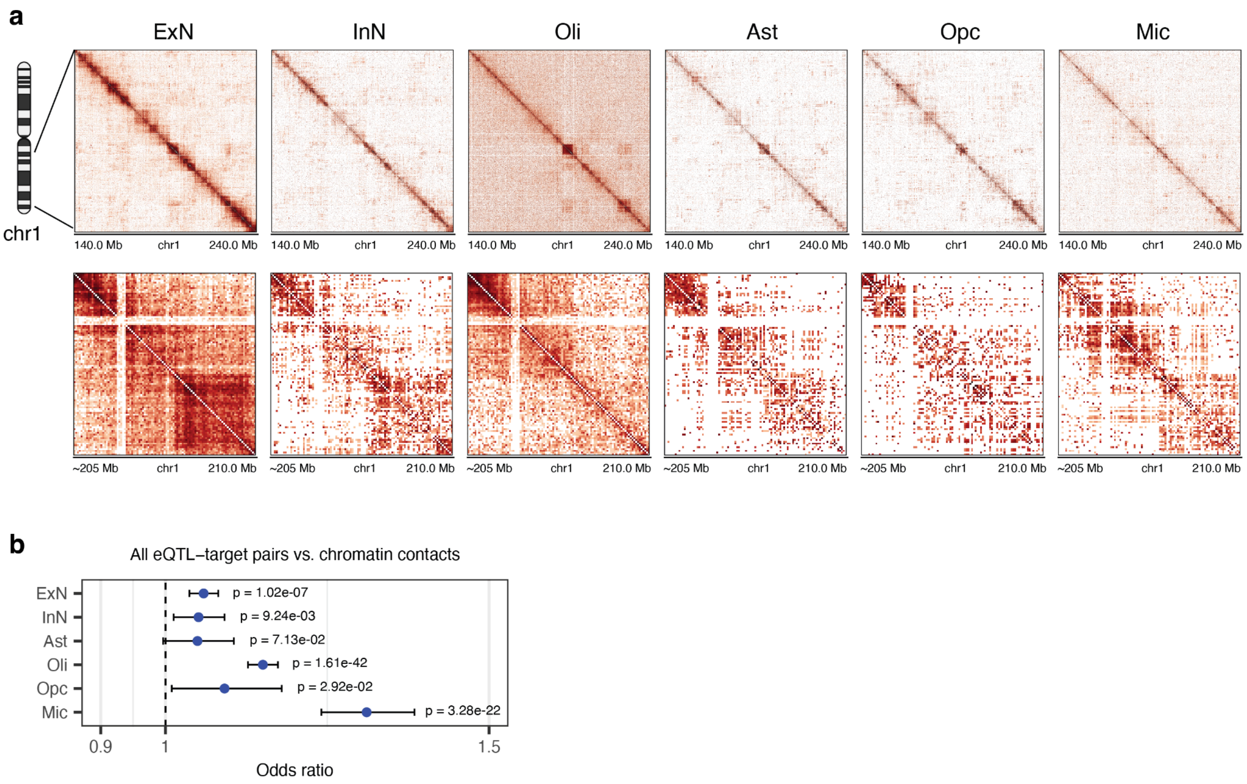


**Supplementary Fig. 3. Cell type specific chromatin contacts and eQTL gene interaction.**

(a) 2-D contact maps for each cortical cell type for chromosome 1 q arm 140Mb to 240Mb region at 1 Mb resolution, and for zoom-in region from 205Mb to 210 Mb at 50 kb resolution. (b) Association tests between the DD contacts and eQTL-target pairs in every cell type, based on all eQTL-target pairs. Odds ratio (center dot) and 95% confidence interval of the odds ratio (whiskers) are plotted for every cell type. One-sided Chi-square test. N = 202,307 DD contacts.
